# R-type voltage-gated Ca^2+^ channels mediate A-type K^+^ current regulation of synaptic input in hippocampal dendrites

**DOI:** 10.1101/2020.05.27.119305

**Authors:** Jonathan G. Murphy, Jakob J. Gutzmann, Lin Lin, Jiahua Hu, Ronald S. Petralia, Ya-Xian Wang, Dax A. Hoffman

## Abstract

The transient K^+^ current (I_A_) carried by pore forming Kv4.2 subunits regulates the propagation of synaptic input, dendritic excitability, and synaptic plasticity in CA1 pyramidal neuron dendrites of the hippocampus. We report that the Ca^2+^ channel subunit Cav2.3 regulates I_A_ in this cell type. We first identified Cav2.3 as a Kv4.2 interacting protein in a proteomic screen and we confirmed Cav2.3-Kv4.2 complex association using multiple techniques. Functionally, Cav2.3 Ca^2+^-entry increases Kv4.2-mediated whole-cell current due to an increase in Kv4.2 surface expression. Using pharmacology and Cav2.3 knockout mice, Cav2.3 was found to promote whole-cell I_A_ and the increasing gradient of I_A_ in the apical dendrite distal to the neuronal soma. Furthermore, the loss of Cav2.3 function leads to enhancement of synaptic currents and spine Ca^2+^ influx. These results present Cav2.3 and Kv4.2 as integral constituents of an ion channel complex that impacts synaptic function in the hippocampus.

## INTRODUCTION

In neuronal dendrites, voltage-gated ion channels modulate the amplitude, propagation and integration of synaptic input. The voltage-gated K^+^ channel subunit Kv4.2 is highly expressed in the dendrites of hippocampal CA1 pyramidal neurons where it assembles into tetrameric channels that conduct a low threshold-activated transient outward K^+^ current known as A-type current (I_A_) (1–5). Dendritic I_A_ regulates neuronal excitability by conducting K^+^ in response to depolarization to attenuate synaptic input, dampen the magnitude of backpropagating action potentials (bAP), oppose dendritic plateau potentials, and limit the size of glutamate uncaging evoked spine Ca^2+^ entry (5–11). As a consequence, the primary A-type channel expressed in CA1 pyramidal neuron dendrites, Kv4.2, plays an active role in shaping propagation of synaptic input, hippocampal synaptic plasticity, and learning (5, 8, 12–18). Dysregulation of I_A_ has been reported in both animal models and human cases of Alzheimer’s disease, epilepsy, and pain sensitization (19–26). A better understanding of the mechanisms underlying neuronal I_A_ would facilitate the identification of therapeutic targets.

The properties of neuronal I_A_ can only be recapitulated in heterologous systems by expression of auxiliary subunits known as K^+^ channel interacting proteins (KChIPs) and dipeptidyl aminopeptidase-like proteins (DPPs), which have profound effects on Kv4.x subunit expression, stability, and biophysical properties (27–30). KChIPs are small (188-285 aa) Ca^2+^- binding proteins of the neuronal Ca^2+^ sensor (NCS) gene family expressed from four genes (*KCNIP1-4*). KChIPs are highly conserved in their globular core domain (∼70%) that contains N and C lobules, each with two EF-hands (EF) surrounding a deep hydrophobic pocket that cradles the N-terminus of the Kv4 subunit (31–33). Like the other NCS proteins, KChIP EFs 2, 3 and 4 bind divalent metal ions whereas EF1 is degenerated such that it cannot (34). At physiological free Ca^2+^ concentrations in neurons, the Ca^2+^ occupancy at the three Ca^2+^-binding EF-hands is unknown. However, dynamic Ca^2+^ binding in response to transient Ca^2+^ elevations during neuronal activity is a compelling Kv4.x channel feedback mechanism. Ca^2+^ binding to purified KChIP induces global structural changes throughout the protein that may regulate oligomerization and Kv4.x interactions (35–39). The effects of Ca^2+^ on the Kv4.x-KChIP complex has been studied using EF-hand mutations or by altering intracellular free Ca^2+^ concentrations (27, 31, 40–42). However, EF-hand mutations potently reduce KChIP regulation of Kv4.x channel trafficking and function, likely disrupting protein tertiary structure independent of Ca^2+^ (43). Furthermore, patch clamp studies of Kv4.x channel function in various neuronal and non-neuronal cell types by addition of intracellular free Ca^2+^ or Ca^2+^ chelators in the patch pipette have produced variable results that are difficult to interpret (41, 44–46). In a similar effort to test the role of elevated intracellular Ca^2+^ on Kv4.x-KChIP function, we recently reported an increase in Kv4.2 current density using a low affinity Ca^2+^ chelator (HEDTA) in the patch pipette to hold intracellular free Ca^2+^ at ∼10 μM in HEK293 cells (42). This effect was unique to a subset of KChIP isoforms suggesting that cell-type specific KChIP expression may be an under- appreciated aspect of Ca^2+^ regulation of Kv4.x-KChIP complexes.

Over the last decade, canonically voltage-gated Kv4.x channels have become appreciated as targets of intracellular Ca^2+^ via Ca^2+^ binding KChIP subunits. In the cerebellum, Kv4.3-mediated I_A_ is regulated by Ca^2+^ entry through T-type voltage-gated Ca^2+^ channels (Cav3.x) in a KChIP-dependent manner that shifts the availability of I_A_ to more negative membrane potentials and modulates cell excitability in an activity-dependent manner (47–50). In hippocampal CA1 pyramidal neurons, Wang and colleagues reported that Cav2.3 (R-type) voltage-gated Ca^2+^ channels function to attenuate the size of evoked EPSPs by promoting Kv4.2 function (51). Cav2.3 channels are expressed in the dendrites and spines of CA1 hippocampal pyramidal neurons and regulate action potential afterhyperpolarization and afterdepolarization, the magnitude of bAP evoked Ca^2+^ transients, and Ca^2+^ influx in spines and dendrites (52–58). Wang et al. disrupted Cav2.3-mediated EPSP boosting using BAPTA or a pan-KChIP antibody in the patch pipette suggesting Cav2.3 and KChIP are in close proximity (< 50 nm). However, the molecular nature of the interaction, whether Kv4.2-KChIP regulation in hippocampus is specific to Cav2.3 channels, and a mechanistic explanation for Cav2.3 regulation of I_A_ remained unknown. Here we identified a nanoscale protein-protein interaction between Cav2.3 and Kv4.2 that regulates the amplitude of neuronal I_A_ to attenuate spontaneous synaptic input through a hippocampus-specific Kv4.2 surface localization mechanism.

## RESULTS

### Cav2.3 and Kv4.2 voltage-gated ion channel subunits form a protein complex in the rodent hippocampus

Dendritic A-type current (I_A_) in CA1 pyramidal neurons requires pore forming Kv4.2 subunits and KChIP and DPP accessory subunits which were identified nearly two decades ago (27, 28). We used tandem affinity purification (TAP) and protein identification with mass spectrometry (MS) to uncover unknown Kv4.2 interactions in hippocampal neurons (59). As expected, Kv4.2 pulled down Kv4.x and DPP and KChIP auxiliary subunits confirming the specificity of the assay (**SFig. 1A**). In addition to known interactions, we also identified peptides representing the Cav2.3 amino acid sequence (**SFig. 1B,C**). Cav2.3 was the only ion channel identified in this screen. To confirm that Cav2.3 and Kv4.2 form a complex, we assessed their distributions in area CA1 of the hippocampus. CA1 neuropil colocalization of Cav2.3 and Kv4.2 immunofluorescent signal was consistent with previous reports of enrichment in hippocampal dendrites (**Fig. 1A**) (52, 60). To evaluate Cav2.3-Kv4.2 colocalization in dendrites and spines, we transfected cultured hippocampal neurons with Cav2.3-GFP (i), Kv4.2-myc (ii), and mCherry (iii) plasmids (**Fig. 1B**). In addition to colocalization in dendrites, Cav2.3-GFP and Kv4.2-myc were enriched in spines relative to the cytosolic mCherry fluorescent protein (**Fig. 1C,D**). To more precisely assess Cav2.3 and Kv4.2 colocalization in spines, we performed double immunogold electron microscopy. Cav2.3 and Kv4.2 particles were visible near synapses in spines and we found evidence of colocalization near the postsynaptic density (**Fig. 1Ei**) and in the spine head (**Fig. 1Eii**). We further confirmed Cav2.3 and Kv4.2 binding by co-immunoprecipitation of native Kv4.2 after Cav2.3 pulldown in lysates from WT hippocampal tissue, but not in lysates isolated from a previously characterized Cav2.3 knockout (KO) mouse line (58, 61), confirming the specificity of the anti-Cav2.3 antibody (**Fig. 1F**). If Cav2.3 and Kv4.2 physically associate, we reasoned that assembly would result in a decrease in FRAP mobility. HEK293 cells were transfected with either YFP-Cav2.3 and CFP or Kv4.2-CFP and YFP and photobleaching was performed in separate cells (**SFig. 2A**). When expressed separately, YFP-Cav2.3 mobile fraction (66.49 ± 0.01%) was considerably larger than Kv4.2-CFP (56.01 ± 0.09%) (**Fig. 1G,I**). To assess whether Cav2.3 and Kv4.2 could reciprocally regulate FRAP mobile fraction, YFP- Cav2.3 and Kv4.2-CFP were coexpressed and bleached simultaneously (**SFig. 2B**). FRAP mobile fraction decreased for both YFP-Cav2.3 (56.28 ± 0.22%) and Kv4.2-CFP (36.77 ± 0.12%) suggesting that Cav2.3 and Kv4.2 bind in non-neuronal cells independent of auxiliary subunits (**Fig. 1H,I**). We next determined if Cav2.3 regulates Kv4.2 mobility in neuronal spines and dendrites. WT or Cav2.3 KO mouse hippocampal neurons were transfected with Kv4.2- GFP and mCherry (structural marker) to assess Cav2.3 regulation of Kv4.2-GFP mobile fraction within small (∼ 0.5 μm^3^) volumes of dendrite shafts or entire spines (**SFig. 2C,D**). Kv4.2-GFP was more mobile in dendrites (55.96 ± 0.01%) than spines (45.00 ± 0.02%) (**Fig. 1L**), consistent with the diffusion limit imposed by the spine neck and reported Kv4.2 interactions with postsynaptic scaffold proteins including PSD95 (62, 63), SAP97 (64), and A-kinase anchoring protein 79/150 (AKAP79/150) (65). Kv4.2-GFP mobile fraction was increased specifically in dendrite shafts (+9.85 ± 3.21%) as opposed to spines (+6.45 ± 3.21%) of Cav2.3 KO mouse neurons (**Fig. 1J-L**). This may be due to a high concentration of immobile Kv4.2 binding partners in spines relative to dendrites. Our FRAP results in both HEK293 cells and neurons confirm a Cav2.3-Kv4.2 complex in hippocampal dendrites and excitatory synapses. These results demonstrate, for the first time, a physical coupling between Cav2.3 and Kv4.2.

**Figure 1.**
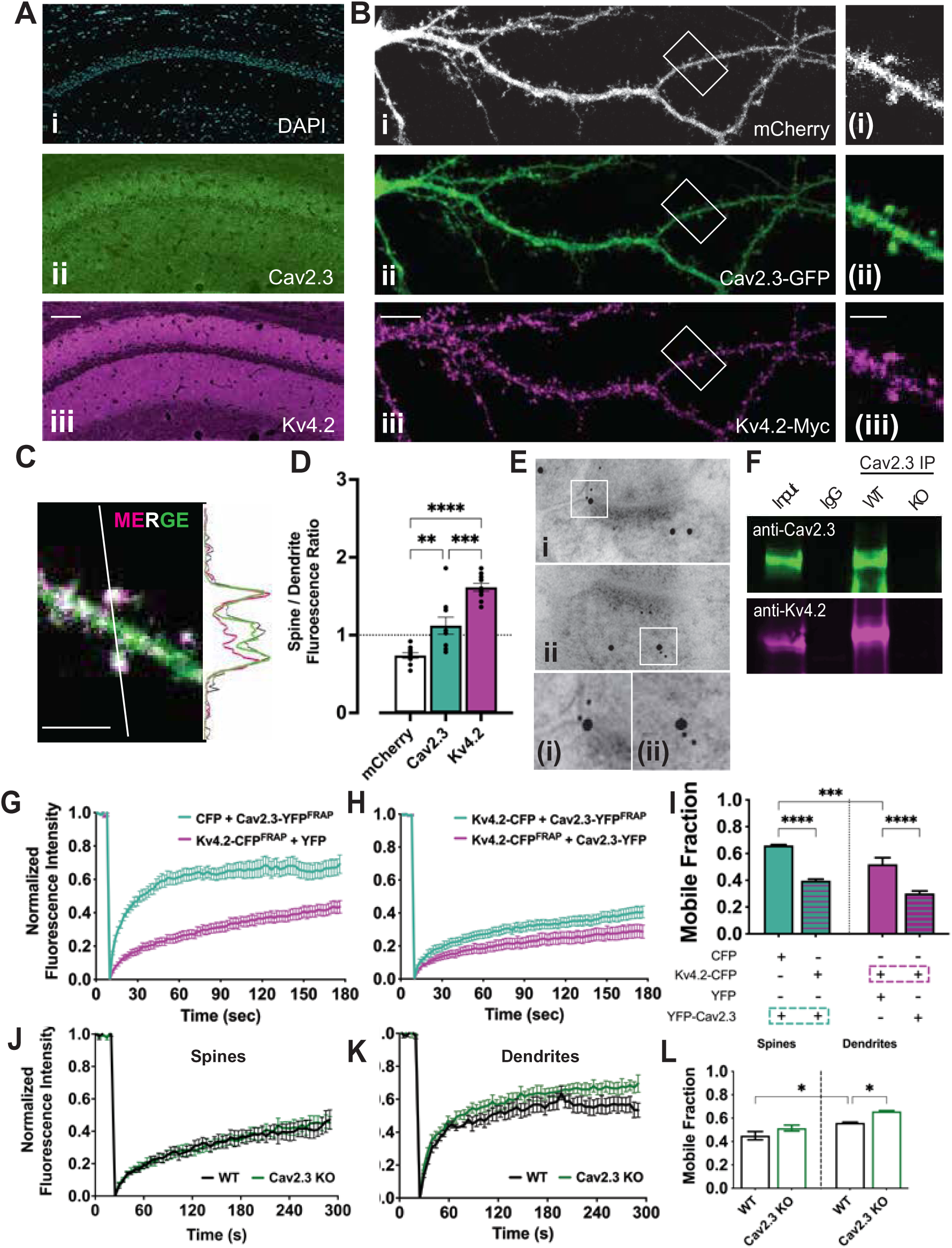
Cav2.3 and Kv4.2 voltage-gated ion channel subunits form a protein complex in the rodent hippocampus. **A.** Mouse hippocampal brain sections were stained for (**Ai**) nuclei (DAPI), (**Aii**) Cav2.3, and (**Aii**) Kv4.2 channels. Cav2.3 and Kv4.2 are localized to dendrite fields of hippocampal area CA1. 100 μm scale bar **B.** Dispersed cultured rat hippocampal neurons expressing (**Ai**, white) mCherry, (**Aii**, green) Cav2.3-GFP, and (**Aiii**, purple) Kv4.2-myc. (Inset) Cav2.3 and Kv4.2 channel fluorescence is enriched in dendritic spines relative to mCherry. 10 μm and 3 μm scale bars **C.** Merged Kv4.2-myc and Cav2.3-GFP fluorescence of dendritic segment in panel B. The representative intensity profile shows enrichment of both Cav2.3-GFP and Kv4.2-myc in spines. 3 μm scale bar. **D.** Cav2.3-GFP and Kv4.2-myc are enriched in spines when compared to cytosolic mCherry. n = 10. **E.** Double immunogold electron micrographs of rat area CA1 hippocampal sections labeled using 5 nm (Cav2.3) and 15 nm gold (Kv4.2). (insets) Cav2.3 and Kv4.2 colocalize within postsynaptic spines. **F.** Cav2.3 immunoprecipitation (green) pulls down Kv4.2 (purple) from native hippocampal tissue. **G.** FRAP recovery curves obtained from HEK293FT cells transfected with either YFP and Kv4.2-CFP (purple) or YFP-Cav2.3 (green) and CFP. Fluorescence recovery within the bleached volume is plotted over time for Kv4.2-CFP and YFP-Cav2.3 when expressed with non-interacting fluorescent proteins (YFP and CFP, respectively). **H.** FRAP recovery curves from HEK293FT cells expressing Kv4.2-CFP and YFP- Cav2.3 after simultaneous CFP/YFP photobleaching. **I.** FRAP recovery curves were fit to exponential functions and the mobile fraction for each transfection condition is plotted. The fluorescent protein species plotted in each bar graph is color-coded with dashed rectangles. Coexpression of Kv4.2-CFP and YFP-Cav2.3 reduces respective mobile fractions consistent with reciprocal interactions between the two channels. (n = 15-19 cells for each condition across 3 experiments). **J.** FRAP recovery curves of spine-localized Kv4.2-SGFP2 expressed in dispersed cultures of WT (black) or Cav2.3 KO (green) mouse hippocampal neurons. **K.** FRAP recovery curves of Kv4.2-SGFP2 localized to dendrite shafts and plotted as in J. **L.** Bar graph comparison of Kv4.2-sGFP2 mobile fraction between WT and Cav2.3 KO mouse neurons. Subcellular comparison showed a significantly greater Kv4.2-SGFP2 mobile fraction in dendrites compared to spines. Kv4.2-SGFP2 mobile fraction was greater in dendrites of Cav2.3 KO mouse neurons when compared to WT. n = 17-18 spines and 14-17 dendrites from 7 WT and 7 Cav2.3 KO neurons. Data was pooled from 2-3 hippocampal cultures. Error bars represent +/- SEM. * *p* < 0.05, ** *p* < 0.01, *** *p* < 0.001, **** *p* < 0.0001. Statistical significance was evaluated by one-way ANOVA with Tukey’s multiple comparisons test.

### Cav2.3 and Kv4.2 bind at a 1:1 channel stoichiometry within a cellular nanodomain

After confirming Cav2.3-Kv4.2 complex formation, we next determined if Cav2.3 and Kv4.2 were in proximity to form a Ca^2+^ nanodomain. FRET is a distance-dependent process that involves the non-radiative transfer of energy from an electronically excited donor to a nearby acceptor fluorophore; FRET decays at the inverse sixth power of the donor and acceptor distance and is detected only within <10 nm (66, 67). To assess Kv4.2 and Cav2.3 proximity, we first introduced Kv4.2-CFP or YFP-Cav2.3 into HEK293 cells with YFP or CFP, respectively, to rule out spurious FRET (0.68 ± 0.34% and 0.46 ± 0.21%) (**Fig. 2A,B**). We then coexpressed Kv4.2-YFP and KChIP3a-CFP, which, as expected, yielded a high FRET efficiency (11.22 ± 0.57%). Coexpression of Kv4.2-CFP and YFP-Cav2.3 resulted in FRET (6.71 ± 0.39%), confirming that they bind within 10 nm (**Fig. 2A,B**). Next, we leveraged the maximum FRET signal at either the donor (*FRET_D,MAX_*) or acceptor (*FRET_A,MAX_*) with saturating concentrations of free acceptors or donors, respectively, to determine the stoichiometry of the Cav2.3-Kv4.2 complex (68). The stoichiometry can be deduced from the maximal efficiency of interacting acceptors and donors expressed as the stoichiometry ratio (*FRET_A,MAX_*/*FRET_D,MAX_*). We validated this method in our hands by taking advantage of known stoichiometries of Kv4.2- KChIP (1:1) and AKAP79-regulatory subunit of protein kinase A, RIIα (1:2) binding. After coexpressing varying ratios of Kv4.2-YFP and KChIP3a-CFP we measured similar maximum FRET efficiencies at both donor and acceptor (*FRET_A,MAX_*: 15.98 ± 0.64%; *FRET_D,MAX_*: 15.75 ± 0.79%) consistent with a 1:1 interaction (**Fig. 2C,F,G**). For AKAP79-YFP and PKA-RII-CFP we measured maximum FRET efficiencies that were most consistent with the expected 1:2 stoichiometry (*FRET_A,MAX_*: 13.28 ± 1.13%; *FRET_D,MAX_*: 5.35 ± 0.26%) (**Fig. 2D,F,G**). Determination of the stoichiometry of YFP-Cav2.3 and Kv4.2-CFP indicated that the FRET complex was most consistent with a 1:4 acceptor:donor ratio that would be expected for a complex containing a single pore-forming YFP-Cav2.3 α1 subunit and a Kv4.2-CFP homotetrameric channel (*FRET_A,MAX_*: 14.52 ± 1.14%; *FRET_D,MAX_*: 3.43 ± 0.33%) (**Fig. 2E,F,G**). These FRET experiments confirm that binding occurs at a scale required for Cav2.3-mediated Ca^2+^ regulation of Kv4.2-KChIP (69, 70).

**Figure 2.**
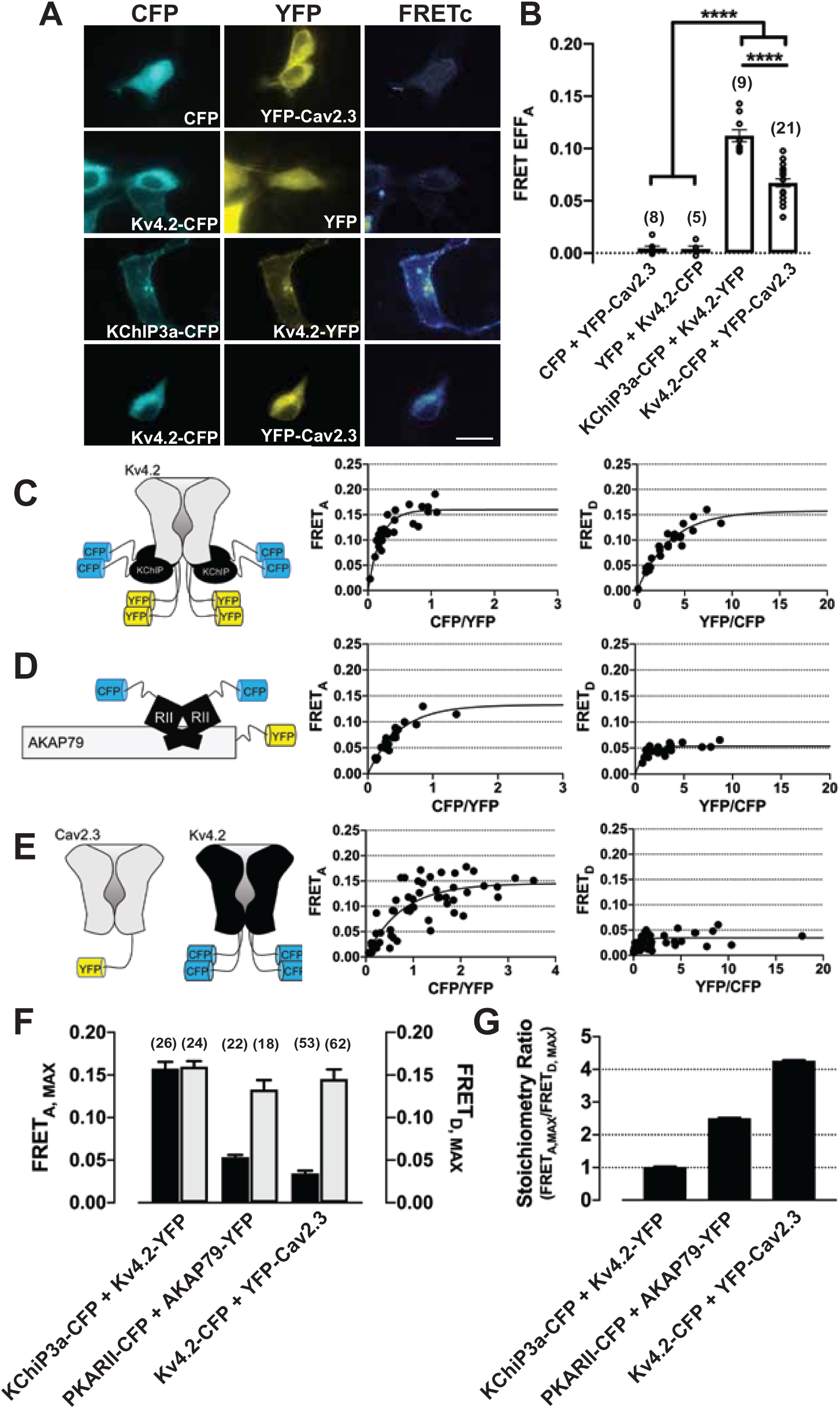
Cav2.3 and Kv4.2 bind at equimolar stoichiometry within a nanodomain. **A.** Representative raw CFP (cyan), YFP (yellow), and FRETc (Temperature spectrum pseudocolor) fluorescent images of HEK293 cells expressing FRET constructs at equimolar concentrations. **B.** Bar graphs show comparison of mean FRET efficiency normalized to acceptor concentration for each condition. Parentheses indicate n values from 2-3 independent experiments. **C-E.** Left, cartoons depict the expected stoichiometry of Kv4.2-YFP and KChIP2c- CFP (1:1) and AKAP79-YFP and PKARIIα-CFP (1:2) while YFP-Cav2.3 and Kv4.2-CFP stoichiometry is inferred from the data. Right, cells were transfected with various ratios of plasmid DNA. Donor and acceptor normalized FRET efficiency were plotted for each cell over the ratio of donor and acceptor fluorescence. **F.** Bar graphs compare acceptor (grey bars) and donor (black bars) normalized FRET efficiency for each condition. Parentheses indicate n values from 2-3 biological replicates. **G.** FRET stoichiometry ratio is calculated using the formula *FRET_A,MAX_*/*FRET_D,MAX_* is plotted for FRET pairs confirming a 1:4 ratio of acceptor:donor or a 1:1 channel stoichiometry due to the tetrameric structure of Kv4.2. Error bars represent +/- SEM.**** *p* < 0.0001. Statistical significance was evaluated by one-way ANOVA with Tukey’s multiple comparisons test.

### Cav2.3 coexpression increases Kv4.2 current density and surface localization in a KChIP- and Ca^2+^-dependent manner

We and others have reported that Kv4.x current density is increased when intracellular free Ca^2+^ is increased to the μM range (42, 71). Nanoscale binding of Cav2.3 and Kv4.2 suggests that Cav2.3 could act as a local μM-source of free Ca^2+^. To test this, we expressed either Kv4.2 alone (**i**), Kv4.2 and KChIP2c (**ii**), or Kv4.2, KChIP2c, and Cav2.3 (**iii**) in HEK293 cells (**Fig. 3A**). As expected, KChIP2c increased Kv4.2 current density and slowed fast inactivation of the macroscopic current (**Fig. 3B,C**). Interestingly, coexpression of Cav2.3 increased Kv4.2 current density (**Fig. 3B,C**) without affecting KChIP-dependent regulation of Kv4.2 voltage-dependence of inactivation and recovery from inactivation (**Fig. 3E,F**). If local Cav2.3-mediated Ca^2+^ influx led to increased Kv4.2 current, this could be disrupted by replacing EGTA with the fast Ca^2+^ chelator BAPTA to limit Ca^2+^ diffusion. BAPTA blocked Cav2.3 increases in Kv4.2 current density (**Fig. 3B,C**) consistent with a Cav2.3-mediated Ca^2+^ influx effect. Coexpression of a Ca^2+^-dead EF-hand mutant KChIP2c, also reversed the Cav2.3 effect (**Fig. 3C**). Increased Kv4.2 current density by Cav2.3 Ca^2+^ influx could be explained by an increase in Kv4.2 surface localization. To test this possibility, we transfected COS7 cells with Kv4.2 alone, Kv4.2 and Cav2.3, Kv4.2 and KChIP2c, or Kv4.2, Cav2.3, and KChIP2c. Kv4.2 surface biotinylation showed that Cav2.3 coexpression led to an increase in Kv4.2 surface localization when compared to KChIP alone (**Fig. 3G,H**). Therefore, assembly of Cav2.3 and Kv4.2 increases Kv4.2 surface localization in a Cav2.3-mediated Ca^2+^ entry and KChIP- dependent manner.

**Figure 3.**
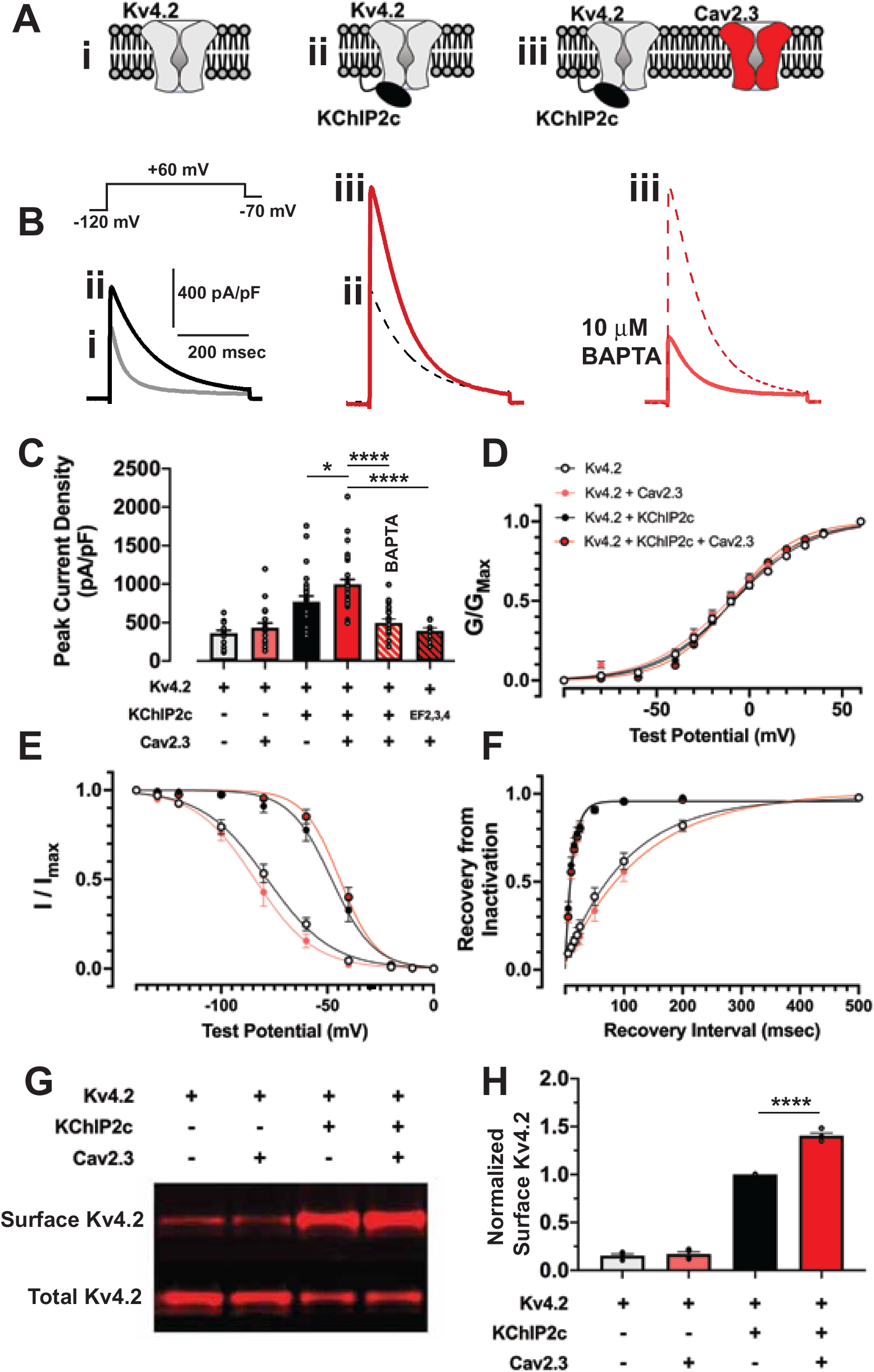
Cav2.3 expression increases Kv4.2 current density in a KChIP- and Ca^2+^- dependent manner in HEK293 cells. **A.** Cartoons depict a subset of transfection conditions with either Kv4.2 alone (**i**), Kv4.2 and KChIP2c (**ii**), or Kv4.2, KChIP2c, and Cav2.3 (**iii**). **B.** Representative traces from conditions shown in A. **C.** Bar graphs plot peak Kv4.2 current density under the conditions shown, n = 10- 32 cells. Cav2.3 expression increases Kv4.2 current density in a KChIP-dependent manner, which is reversed by replacement of EGTA with BAPTA in the patch pipette and by coexpression of EF-dead KChIP2c. **D.** Kv4.2 voltage-dependence of activation is plotted using normalized conductance against a range of membrane test potentials and fit to a Boltzmann function. **E.** Kv4.2 voltage-dependance of inactivation is plotted using normalized current against conditioning test potentials and fit to a Boltzmann function. **F.** Kv4.2 recovery from inactivation is plotted as the fraction of current recovered using a +60 mV test potential from an initial test potential of the same magnitude against various recovery intervals. **G.** Representative western blot of a surface biotinylation assay in COS7 cells transfected according to the indicated conditions. **H.** Bar graph shows Kv4.2 surface expression normalized by day of experiment to the Kv4.2 and KChIP2c expression condition. Error bars represent +/- SEM from 4 biological replicates. \**p* < 0.05, \*\*\*\**p*< 0.0001. Statistical significance was evaluated by one-way ANOVA with Tukey’s multiple comparisons test.

### Cav2.3 promotes I_A_ in cultured hippocampal neurons

After we established functional regulation of Kv4.2 by Cav2.3 in HEK293 cells, we wanted to confirm Cav2.3 regulation of native neuronal I_A_. Whole-cell I_A_ density in cultured hippocampal neurons reaches a maximum at 6-9 days in vitro (DIV) prior to synaptic maturation (**SFig. 3A,B**). The peak outward K^+^ current was enriched for I_A_ relative to the sustained current (92.49 ± 0.02%; data not shown); therefore, we monitored total peak outward current as a proxy for I_A_. Voltage clamp recordings were stable over a 360 s recording period (**Fig. 4Ai,B,C**). We confirmed the presence of Kv4.x containing channels using 500 nM AmmTx3, a Kv4.x selective toxin (**Fig. 4Aiii,B,C**) (72, 73). Ni^2+^, a selective Cav2.3 blocker, was used as an alternative to the more potent and selective Cav2.3 toxin SNX-482 because SNX-482 has been reported to potently inhibit Kv4.x channels (74–76). Application of 200 μM Ni^2+^ led to a rapid and sustained reduction in I_A_ (**Fig. 4Aii,B,C**). Ni^2+^ regulation of I_A_ was dose-dependent and the apparent IC_50_ (∼44.1 μM) approximated values reported for Cav2.3 (27.4-66.0 μM) (**SFig. 4A**) (77). Ni^2+^ did not affect voltage dependence of inactivation for I_A_, suggesting a distinct mechanism in hippocampus relative to cerebellum (**SFig. 4B**) (47–50). Cav2.3 regulation of Kv4.2 function may involve surface expression as observed above in HEK293 cells. To determine if Ni^2+^ block of Cav2.3 channels mediated a reduction in surface Kv4.2 channels, we performed Kv4.2 surface biotinylation in cultured hippocampal neurons treated with 200 μM Ni^2+^. We detected a reduction in Kv4.2 surface expression in treated neurons when compared to untreated controls (**Fig. 4D**). We repeated time courses of whole-cell I_A_ using selective antagonists of other known neuronal VGCCs to determine if the effects of Ni^2+^ were specific to Cav2.3. Blockers for T-type (5 μM TTA-P2), L-type (5 μM Nimodipine), and P/Q or N-type (1 μM ω-conotoxin GVIA + 3 μM ω- conotoxin MVIIC) did not reduce whole-cell I_A_ (**Fig. 4E**). Ni^2+^ led to a larger reduction in whole-cell I_A_ in WT vs. Cav2.3 KO neurons (72.11 ± 0.04% vs. 81.51 ± 0.01%, *p = 0.049*). Taken together, Ni^2+^ action on Cav2.3 channels acutely regulates Kv4.2 surface localization in hippocampal neurons.

**Figure 4.**
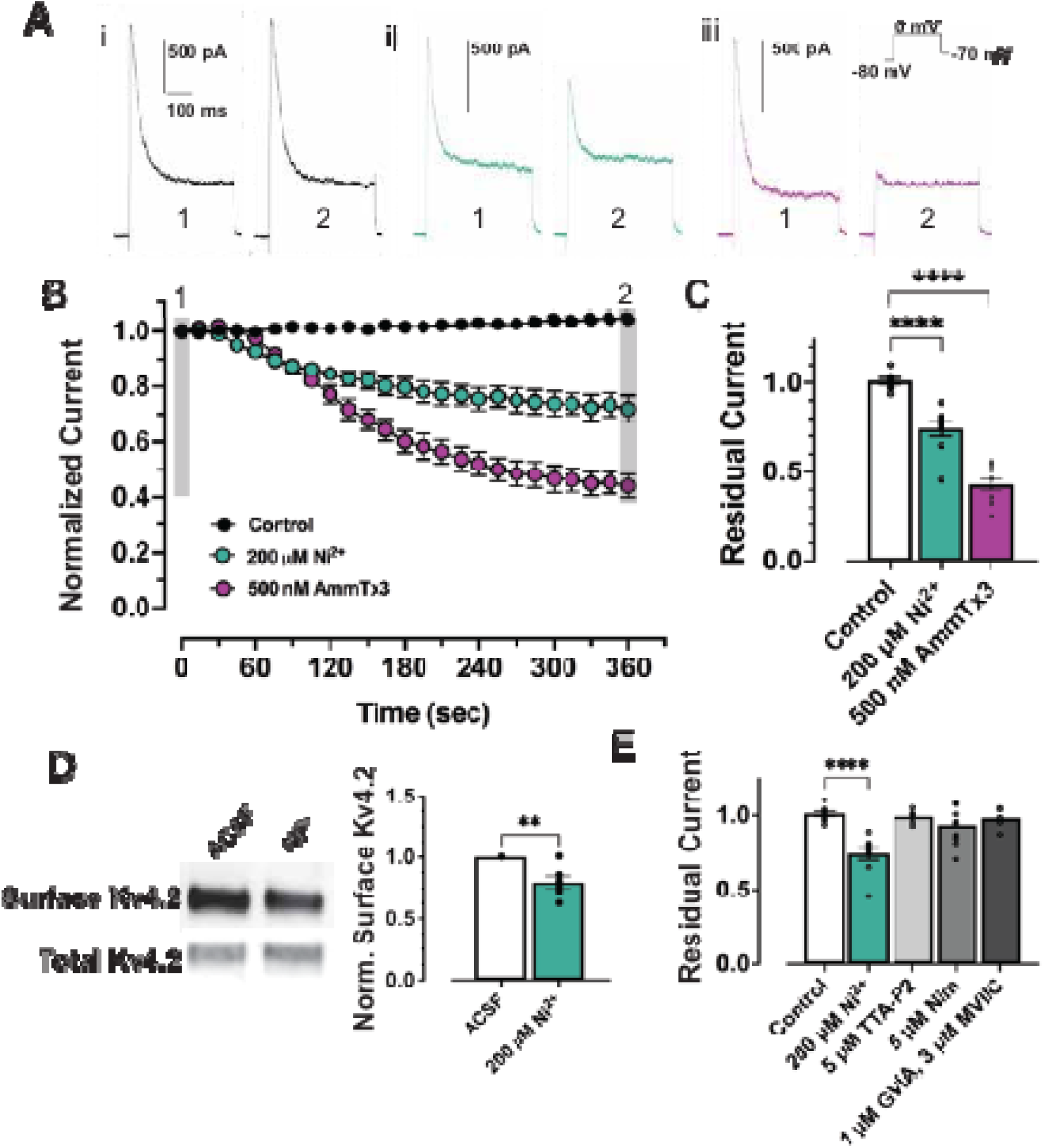
Cav2.3 promotes I_A_ in cultured hippocampal neurons. **A.** Representative whole-cell voltage clamp currents recorded from DIV7-9 neurons grown in dispersed rat hippocampal cultures. After obtaining whole-cell configuration, neurons were either (i) untreated and superfused with ACSF, or rapidly exchanged into ACSF containing (ii) 200 μM Ni^2+^ or (iii) 500 nM AmmTx3. First (1) and last (2) traces in the 6 min time course are shown. **B.** Averaged peak outward current normalized to current at break-in is plotted against the duration of the experiment. Drug wash-in began at 15 s. **C.** Peak residual current amplitude remaining at the end of the recording period was normalized to current at break-in and plotted for each condition. **D.** Representative western blot of a surface biotinylation assay in dispersed rat hippocampal cultures. Endogenous Kv4.2 surface expression was reduced following a 10 min incubation in ACSF supplemented with 200 μM Ni^2+^ when compared to non-treated neurons. **E.** Residual transient current is plotted after treatments with neuronal T-type (5 μM TTA-P2), L-type (5 μM Nimodipine) and P/Q and N-type (1 μM ω-conotoxin GVIA / 5 μM ω- conotoxin MVIIC) blockers. Note data from panel C was reused for comparison. Statistical significance was evaluated by either student’s t-test or one-way ANOVA with Tukey’s multiple comparisons test. * *p* < 0.05, ** *p* < 0.01, *** *p* < 0.001, **** *p* < 0.0001.

### The I_A_ gradient is disrupted in Cav2.3 KO CA1 pyramidal neuron dendrites

Kv4.2 channels in CA1 neurons regulate active processes in dendrites to transform synaptic input and set the threshold for synaptic plasticity. Therefore, we wanted to determine the state of I_A_ in CA1 dendrites of Cav2.3 KO mice. Somatic I_A_ was reduced in whole-cell recordings from Cav2.3 KOs when compared to WT (**Fig. 5A**). This could be explained by a reduction in I_A_-conducting channels at the somatic cell membrane. However, outside-out patches pulled from the cell soma did not replicate the whole-cell reduction in I_A_ (**Fig. 5B**). We next asked if the whole-cell reduction in I_A_ observed in Cav2.3 KO recordings arose from a specific loss of dendritic Kv4.2 channels. To determine this, we performed cell-attached dendritic recordings along the apical dendrite (**Fig. 5C**). Comparison of distal versus proximal I_A_ suggested that the reduced I_A_ measured at the soma is due to a reduction in dendritic I_A_ conducting channels in Cav2.3 KO neurons (**Fig. 5D**). These results suggest that Cav2.3 channel expression regulates the characteristic graded expression of I_A_ in dendrites.

**Figure 5.**
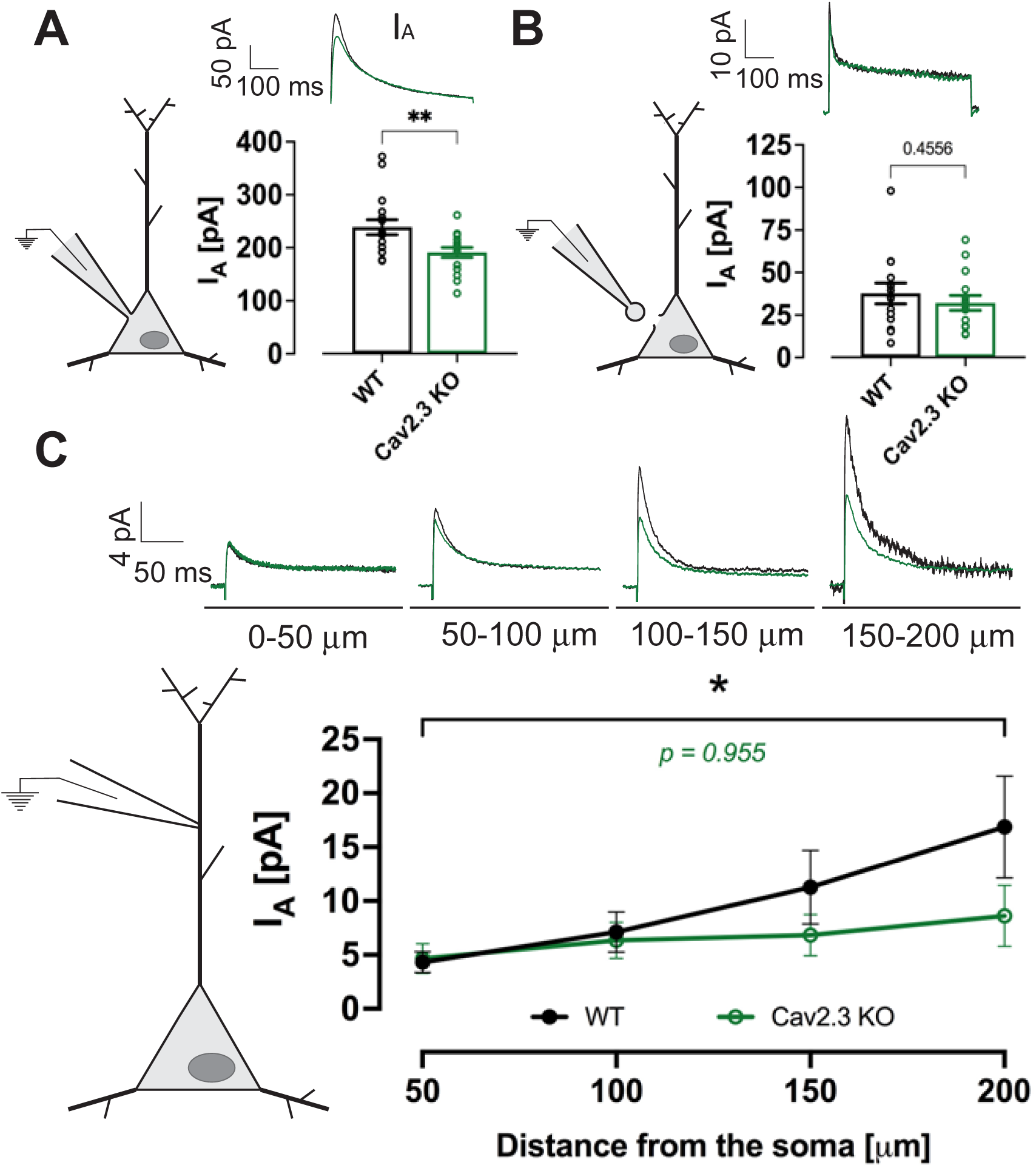
The I_A_ gradient is disrupted in Cav2.3 KO CA1 pyramidal neuron dendrites. **A.** Comparison of whole-cell I_A_ amplitude in WT (black) and Cav2.3 KO (green) mouse; n = 17 WT and 16 KO cells. Representative I_A_ recordings for WT and Cav2.3 KO are shown. **B.** Comparison of I_A_ in outside-out somatic patches of WT and Cav2.3 KO; n = 14 WT and 15 KO cells. Representative I_A_ recordings are shown using averaged data. **C.** (Above) Representative cell-attached I_A_ recordings from the apical dendrites of WT and Cav2.3 KO neurons at various distances from the neuronal cell body. (Below) I_A_ is plotted for dendrite segments. The I_A_ gradient is significantly reduced in Cav2.3 KO dendrites relative to WT. Error bars represent +/- SEM. * *p* < 0.05, ** *p* < 0.01. Statistical significance was evaluated by either student’s t-test or one-way ANOVA with Tukey’s multiple comparisons test.

### Cav2.3 promotes I_A_-mediated attenuation of spontaneous synaptic currents

Voltage-gated ion channels near excitatory synapses open in response to glutamate receptor-mediated depolarization (9, 56, 78, 79). As a consequence, the amplitude of the synaptic current measured at the cell soma represents the integration of glutamate receptor and voltage-activated membrane conductances. Kv4.2 and Cav2.3 channels are localized to CA1 spines and dendrites where they affect the synaptic potential and shape its propagation (5, 51, 52, 56, 80–82). We have previously shown that I_A_ reduces the magnitude of spontaneous miniature EPSCs (83). Therefore, if Cav2.3 channels boost I_A_, Ni^2+^-mediated reduction in I_A_ could enhance mEPSCs. To test this, we applied Ni^2+^ and recorded spontaneous mEPSCs (**Fig. 6A**). Ni^2+^ produced a rightward shift in the distribution of mEPSC amplitudes, though with smaller effect than blockade of Kv4.2 with AmmTx3 (**Fig. 6B**). Ni^2+^ and AmmTx3 together mimicked AmmTx3 alone, suggesting that Kv4.2 channel block occluded the Ni^2+^ effect without any synergism (**Fig. 6B,C**). Despite an increase on mEPSC amplitude, Ni^2+^ reduced mEPSC frequency. While outside the scope of this study, this might suggest a presynaptic effect of Ni^2+^ on VGCCs, including T-type, that control glutamate release in the hippocampus (**Fig. 6C**) (51, 74, 84). To assess if Cav2.3 expression was required for Kv4.2 regulation we measured the effect of AmmTx3 on mEPSCs in WT or Cav2.3 KO mouse neurons. mEPSCs recorded from WT neurons treated with AmmTx3 were larger in amplitude than untreated controls (**Fig. 6D-F**). As expected, AmmTx3 did not increase mEPSC amplitudes in Cav2.3 KO mouse neurons (**Fig. 6G-I**). Kv4.2 channels are also reported to be clustered at GABAergic synapses where they may regulate inhibitory synaptic input (85–88). Using patch electrodes containing high Cl^-^, we measured GABAergic mIPSCs in WT mouse neurons at a holding potential of -70mV to optimize availability of Kv4.2 channels (**SFig. 5A**). AmmTx3 increased mIPSC amplitude in WT but not Cav2.3 KO mouse neurons (**SFig. 5B-F**). The above results support a role for Cav2.3 as an integral component of Kv4.2 channel complexes required for regulation of synaptic input.

**Figure 6.**
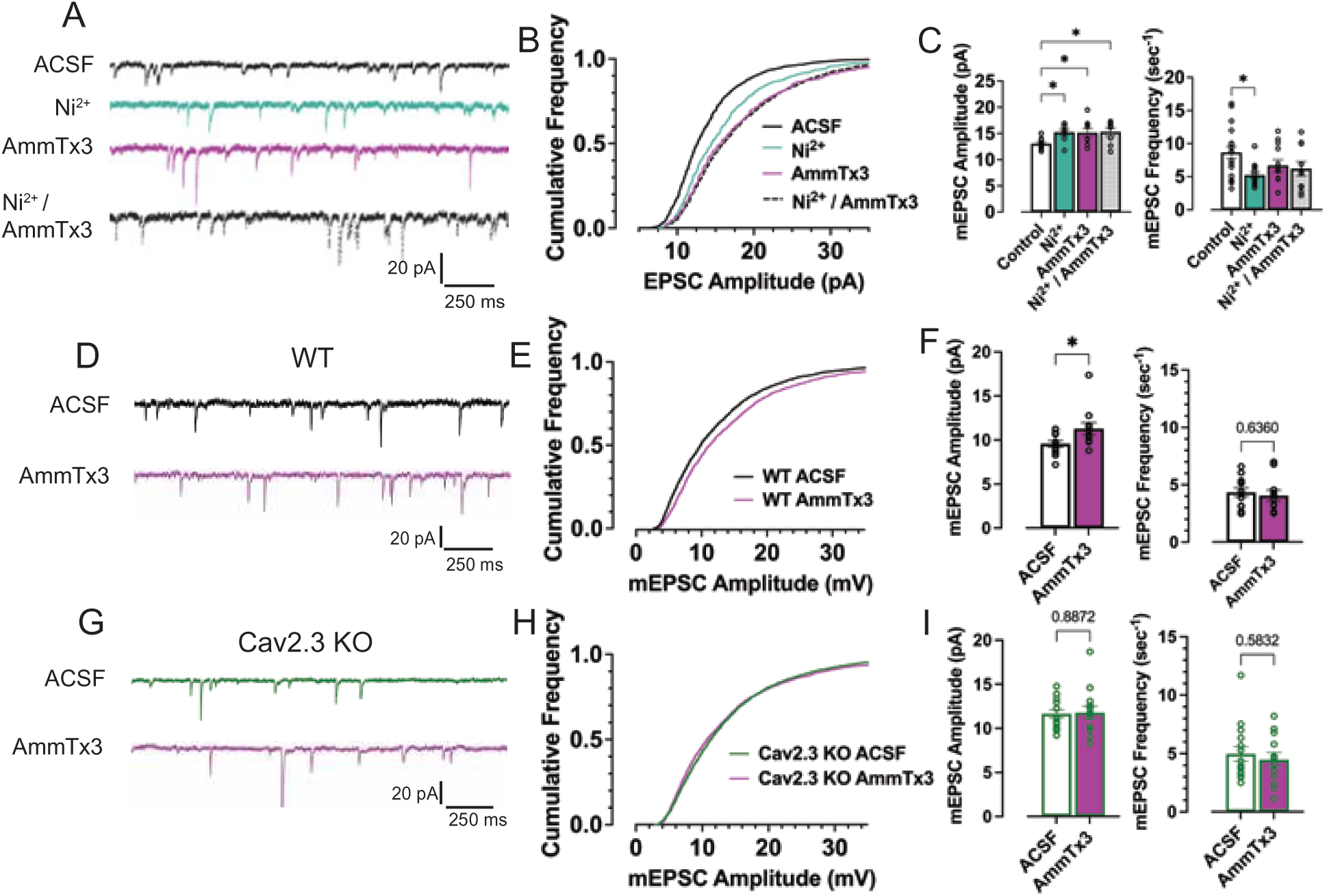
Cav2.3 promotes I_A_-mediated attenuation of spontaneous miniature excitatory postsynaptic currents. **A.** Representative miniature excitatory postsynaptic currents (mEPSCs) recorded from dispersed rat hippocampal cultures at a holding potential of -70 mV in the absence (black trace) or presence of Ni^2+^ (green), AmmTx3 (purple) or Ni2+ and AmmTx3 combined (hashed trace). The cumulative distribution of mEPSC amplitudes. Distribution of mEPSCs in the presence of Ni^2+^, AmmTx3, or Ni^2+^ and AmmTx3 in combination are shifted toward a higher proportion of larger events. **C.** Mean of the Median event amplitude (left) and mean frequency (right) from each neuron were compared. AmmTx3, Ni^2+^, and Ni^2+^/AmmTx3 increases median amplitude. However, there is no additive effect of Ni^2+^ and AmmTx3. Mean mEPSC frequency is decreased in the presence of Ni^2+^. **D.** Representative mEPSCs recorded from WT mouse neurons in the absence (black trace) or presence (purple trace) of AmmTx3. **E.** Cumulative frequency distribution of WT mEPSCs. AmmTx3 induces a rightward shift in mEPSC amplitudes. **F.** Bar graphs are plotted as in C. AmmTx3 increases mEPSC amplitudes as in WT mouse. **G-I.** Cav2.3 KO mEPSCs recorded and summarized as in D-F. AmmTx3 had no effect on mEPSC amplitude in Cav2.3 KO neurons. Error bars represent +/- SEM. * *p* < 0.05. Statistical significance was evaluated by unpaired t-test or one-way ANOVA with Tukey’s multiple comparisons test.

### Cav2.3 promotes I_A_-mediated attenuation of spontaneous quantal spine Ca^2+^ signals

If the Cav2.3-Kv4.2 complex regulates synaptic currents recorded at the cell soma as shown above and by Wang et al. 2014, then it is possible that quantal spontaneous currents in spines may be similarly affected. Fluorescent Ca^2+^ indicators are suited for visualization of NMDAR-mediated Ca^2+^ entry in response to spontaneous quantal glutamate release events (89–91). We transfected neurons with the fluorescent Ca^2+^ indicator GCaMP6f and mCherry as a cell-filling marker. The frequency, amplitude, and duration of NMDAR-mediated Ca^2+^ signals were highly variable both among and within individual spines (**Figure 7A**). The NMDAR blocker AP5 abolished Ca^2+^ signals consistent with a requirement for NMDARs Ca^2+^ influx (**Fig. 7A,B**). AmmTx3 block of Kv4.2 channels did not alter the amplitude of Ca^2+^ signals but did lengthen the event half-width suggesting that Kv4.2 channels oppose spine depolarization initiated by glutamate receptors (**Fig. 7C**). Kv4.2 block increased the integrated amount of Ca^2+^ entry (**Fig. 7D**). As expected, Ni^2+^ reduced integrated spine Ca^2+^ signals possibly due to a combinatorial effect on Ni^2+^ sensitive pre- and postsynaptic VGCCs (**Fig. 7E**). To determine if Cav2.3 Ca^2+^ entry modulated the function of Kv4.2 channels, we preapplied Ni^2+^ followed by coapplication of AmmTx3. Preapplication of Ni^2+^ occluded the AmmTx3-mediated increase in spine Ca^2+^ suggesting Cav2.3 channels are required to maintain spine repolarization by Kv4.2 channels (**Fig. 7F**). Unlike WT mouse neurons, we found that quantal spine Ca^2+^ signals were not amplified in Cav2.3 KO mouse neurons treated with AmmTx3 (**Fig. 7G,H**). Loss of Kv4.2 regulation led to larger basal spine Ca^2+^ signals in Cav2.3 KO neurons when compared to WT (**Fig. 7I**). In consequence, Cav2.3 function and expression is required for Kv4.2-mediated regulation of spontaneous quantal spine Ca^2+^ events.

**Figure 7.**
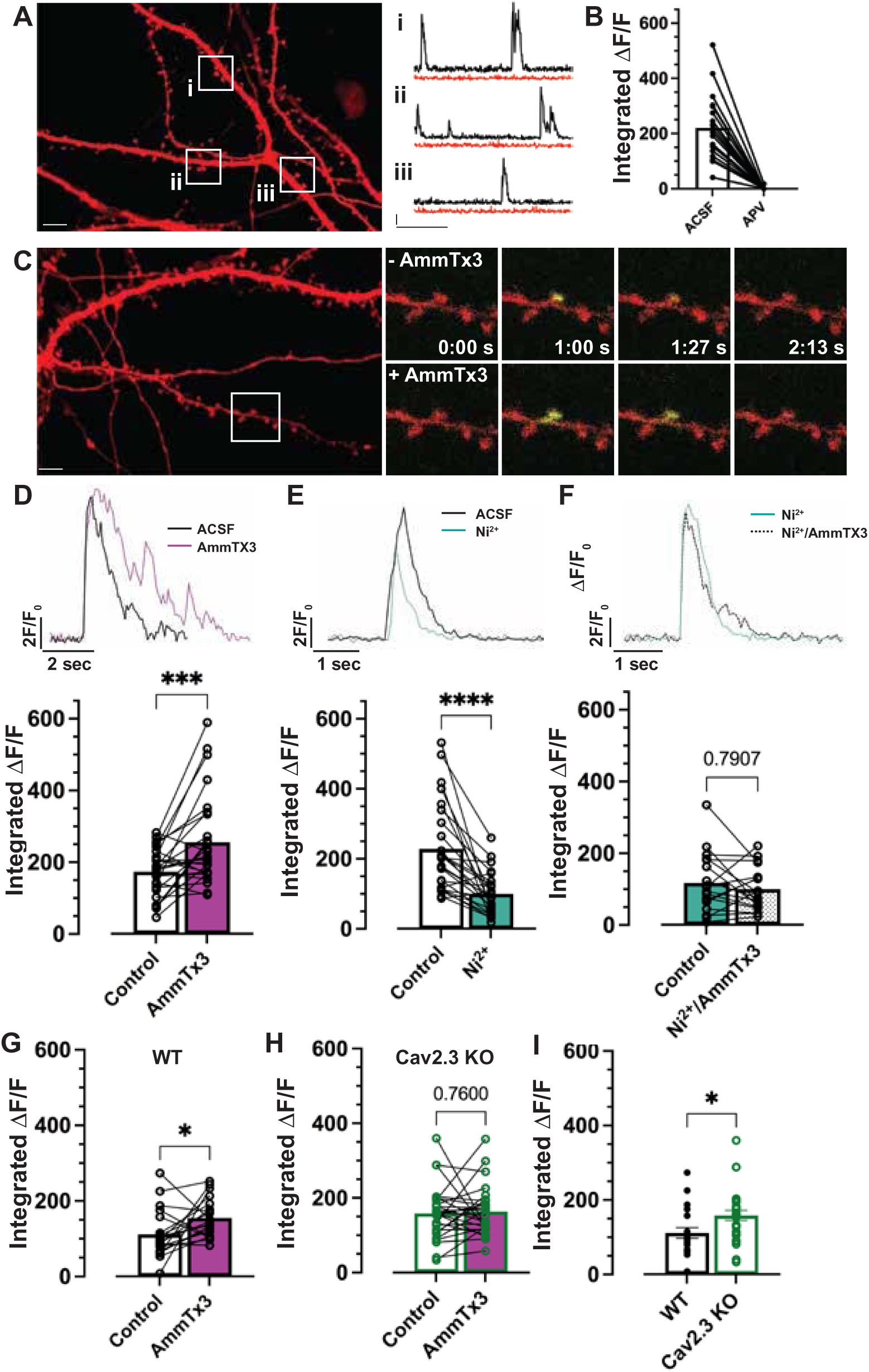
Cav2.3 promotes I_A_-mediated attenuation of spontaneous quantal spine Ca^2+^ signals. **A.** (left) A representative cultured rat hippocampal neuron expressing mCherry (red) and GCaMP6f (not shown). (right) Spontaneous spine Ca^2+^ transients in three representative spines (i, ii, and iii). Traces plot the magnitude of GCaMP6f fluorescence (ΔF/F_0_) over time for each representative spine before (black traces) and after a 10 min application of D-AP5 (50 μM) (red traces). Scale bar: y = 2 ΔF/F_0_; x = 10 s. **B.** The integral of each Ca^2+^ transient for many spines is plotted before and after application of D-AP5. Spine Ca^2+^ signals require the function of NMDARs. **C.** Left, A neuron transfected as in A, but treated with AmmTx3 (500 nM). Right, time lapse images of a representative spine before (above) and after (below) AmmTx3 treatment. **D-F.** (above) The magnitude of GCaMP6f fluorescence (ΔF/F_0_) over time in a representative spine is plotted before and after the indicated drug treatment. (below) Integrated Ca^2+^ influx is plotted for many spines before and after the indicated conditions. AmmTx3 enhanced and Ni^2+^ decreased spine Ca^2+^ signals. Ni^2+^ pretreatment blocked AmmTx3-mediated enhancement of spine Ca^2+^ influx. **G.** AmmTx3 treatment increased spine Ca^2+^ influx in WT neurons. **H.** AmmTx3 does not enhance spine Ca^2+^ signals in Cav2.3 KO neurons. **I.** Basal spine Ca^2+^ transients are increased in Cav2.3 KO neurons. * *p* < 0.05, *** *p* < 0.001, ****p < 0.0001. Statistical significance was evaluated by paired or unpaired t-tests.

## DISCUSSION

The present study was designed to both confirm and define the nature of a Cav2.3- Kv4.2 complex first identified in a proteomic screen and to assess its functional role in pyramidal neurons of the hippocampus. We showed that Cav2.3 and Kv4.2 colocalize at 1:1 stoichiometry within dendritic spine nanodomains independent of auxiliary subunits (**Figures 1,2**). Functionally, we found that the Cav2.3-Kv4.2 interaction increases Kv4.2 surface localization and contributes to maintenance of the I_A_ dendritic gradient (**Figures 3-5**). Previous studies have demonstrated that dendritic I_A_ regulates excitability, spine Ca^2+^ entry, back propagation of action potentials, plateau potentials, synaptic plasticity, and hippocampus-dependent learning (92–94). By regulating the magnitude of I_A_, the Cav2.3-Kv4.2 complex therefore plays a fundamental role in dendritic function. Cav2.3-mediated Ca^2+^ entry likely promotes the reported roles for dendritic I_A_. Indeed, we found that Cav2.3 channels are required for AmmTx3-mediated boosting of mEPSCs, mIPSCs, and spine Ca^2+^ signals (**Figures 6,7**).

The proximity of the Cav2.3-Kv4.2 interaction is critical for regulation of synaptic input. Our FRET data suggests that Cav2.3 and Kv4.2 bind directly since we estimate the diameter of the Kv4.2-KChIP complex itself to be close to ∼10 nm (the upper limit of FRET) based on the Kv2.1-β_2_ channel structure (95). Therefore, KChIPs arrayed around the intracellular T1 domain of the Kv4.2 channel could be exposed to Ca^2+^ concentrations approaching 10 μM during Cav2.3 channel openings based on Ca^2+^ source diffusion models (70). While the steady-state Ca^2+^ occupancy of KChIP is unknown *in vivo*, dynamic KChIP Ca^2+^ binding may account for Cav2.3-mediated regulation of Kv4.2 surface expression through reduced channel turnover. In a test tube, Ca^2+^ binds to KChIP at high (EF3 and EF4) and low affinity sites (EF2) and Ca^2+^ binding promotes KChIP folding and binding to Kv4.x channels (31, 36, 37, 43). While these studies are informative, an *in vivo* mechanism to explain how dynamic KChIP-Ca^2+^ exchange may regulate Kv4.x function remains elusive. In the present study, we report measurements of I_A_ in cultured neurons and hippocampal slices to provide evidence for Cav2.3 regulation of the magnitude of I_A_ (**Figures 4,5**). Cav2.3-mediated Ca^2+^ entry was required to increase Kv4.2 functional expression in a KChIP-dependent manner in non-neuronal cells (**Figure 3**). Several lines of evidence support a Cav2.3-mediated Ca^2+^- and KChIP-dependent surface expression mechanism as opposed to regulation of Kv4.2 conductance or gating in CA1 pyramidal neurons. First, increased single channel conductance has ostensibly been ruled out in previous studies as a mechanism for the increase in Kv4.2 current density mediated by KChIPs (96, 97). However, it is yet to be determined if elevations of intracellular Ca^2+^ may regulate Kv4.2 single channel conductance. Second, we reported previously that elevated intracellular Ca^2+^ led to an increase in Kv4.x current density without affecting classical KChIP-dependent processes including inactivation gating (42). However, stoichiometric KChIP binding is expression dependent (98–100) and if Ca^2+^ shifted Kv4.x binding affinity for KChIP this might be overcome by the overexpression of KChIP relative to Kv4.x in those studies (∼8:1 molar excess). Lastly, we found increased Kv4.2 surface localization by Cav2.3 coexpression in non-neuronal cells and Ni^2+^ reduced Kv4.2 surface localization in neurons in surface biotinylation experiments. Taken together, Cav2.3 and Kv4.2 functional coupling in hippocampal neurons increases the Kv4.2 current amplitude through enhanced surface expression or stability. Therefore, it would seem that increased surface localization of Kv4.2 channels is the likely mechanism to explain the reported EPSP boosting by Cav2.3 block (51). This is in contrast to the mechanism reported for the cerebellar Cav3.x-Kv4.3 complex that promotes I_A_ via a shift in channel availability to more negative potentials (47, 48). It remains to be determined what mediates the disparate regulatory mechanisms in hippocampus and cerebellum. We demonstrated the specificity of Cav2.3 regulation of neuronal I_A_ in hippocampus using VGCC pharmacology and Cav2.3 KO mice (**Figure 4E**). Our observation of residual whole-cell I_A_ regulation by Ni^2+^ treatment in DIV 6-9 Cav2.3 KO neurons, though less so than in WT, suggests there may be compensatory mechanisms to maintain I_A_ in the absence of Cav2.3 expression (**Figure 4F,G**). Perhaps other VGCC subtypes may be able to regulate Kv4.x channels in the absence of Cav2.3, particularly in immature neurons with few synapses. Importantly, compensation did not salvage the dendritic I_A_ gradient nor loss of I_A_-mediated attenuation of synaptic input in Cav2.3 KO adult mouse hippocampal neurons (**Figures 5-7**).

This is the first report describing voltage-gated channel-mediated regulation of quantal NMDAR-mediated Ca^2+^ signals (**Figure 7**). In prior studies, pharmacological block of either AMPAR or voltage-gated channels had no measurable effect on quantal NMDAR-mediated Ca^2+^ transients (101, 102). However, subthreshold synaptic depolarizations can activate voltage-gated channels including R- and T-type Ca^2+^, and A-type K^+^ channels, all of which are activated at relatively hyperpolarized membrane potentials (5, 9, 56, 78, 79). Why did prior studies not find a contribution of voltage-gated channels to the quantal NMDAR-mediated Ca^2+^ signals? The Ca^2+^ signal variability between any two spines necessitated the more straightforward approach of within-spine comparisons. We also found that AmmTx3 regulated the duration of spine Ca^2+^ events more so than amplitude, leading us to compare Ca^2+^ signal integrals. Kv4.2 regulation of NMDAR Ca^2+^ signals is consistent with the known role of Kv4.2 channels in synaptic plasticity and implicates Cav2.3 as a critical component of an ion channel complex that regulates NMDAR-mediated synaptic signaling.

I_A_ increases linearly ∼5-fold on the distal apical dendrites 350 μm from the soma of rat CA1 pyramidal neurons (5). Curiously, histochemical measurements of Kv4.2 expression levels suggest at most a 2-fold increase in expression along the proximal-distal extent of the stratum radiatum (1, 60, 80, 103). A constellation of studies supports both auxiliary subunit and enzyme-dependent pathways may underlie the disparity between Kv4.2 expression and function. The DPP6 auxiliary subunit establishes the Kv4.2 functional gradient in CA1 pyramidal neurons through enhanced dendrite-directed surface expression and increased availability of Kv4.2 at hyperpolarized membrane potentials (15). Distal dendritic I_A_ activates at more negative membrane potentials when compared to proximal or somatic I_A_ (5), and this is reversed by activity of PKA, protein kinase C (PKC), or mitogen-activated kinase pathways (12, 104, 105). During induction of synaptic plasticity, Kv4.2 channels internalize in an NMDAR- and Ca^2+^- dependent manner by PKA phosphorylation at Ser522 (63, 83). Furthermore, Kv4.2 surface expression is bidirectionally regulated by local signaling of the Ca^2+-^activated phosphatase calcineurin (CaN) and PKA through the postsynaptic scaffolding protein AKAP79/150 (65). Another Ca^2+^-dependent enzyme, Ca^2+^-calmodulin dependent kinase II, has also been shown to promote Kv4.2 surface expression through direct channel phosphorylation (106). We recently reported a role for a proline isomerase, Pin1 in activity-dependent downregulation of Kv4.2 function downstream of p38 phosphorylation of T607 (59). Thus, differential localization of kinases and phosphatases along the proximal-distal axis of CA1 pyramidal neurons could underlie the I_A_ gradient. It is possible that Cav2.3 Ca^2+^ entry may impinge on these pathways in a Ca^2+^-nanodomain fashion. For example, AKAP79/150 localizes PKA and CaN for phospho-regulation of AMPAR and L-type Ca^2+^ channels (107). Hypothetically, Cav2.3 Ca^2+^ entry could also drive activation of AKAP-anchored CaN and dephosphorylate Kv4.2 channels, favoring surface expression. Deciphering the mechanistic link between local Ca^2+^ influx and increased Kv4.2 surface expression is a goal for future research.

KChIP heterogeneity is a largely unexplored area of research in CA1 pyramidal neurons. Alternative splicing and variable start codons of the *KCNIP1-4* transcripts result in up to 17 distinct KChIP isoforms with unique N-terminal domains. Variant KChIPs have been categorized based on the tendency for the N-terminus to mediate membrane interactions (33). A subset of “cytoplasmic” KChIP isoforms confer Ca^2+^ regulation onto Kv4.2 channel complexes, N-myristoylated isoforms do not, and reversibly palmitoylated isoforms have mixed Ca^2+^ sensitivity (42). Members of a fourth transmembrane class of KChIPs retain Kv4.2 complexes within the ER and reduce Kv4.2 surface expression when compared with other KChIP isoforms (108). In hippocampus, there is overlapping expression of both Ca^2+^ sensitive and Ca^2+^ insensitive KChIP isoforms. Could differential subcellular targeting of KChIPs lead to increased Kv4.2 function in distal dendrites? Perhaps membrane localized KChIPs, like the Kv4.2 suppressor KChIP4a, may be restricted to soma and proximal dendrites. *KCNIP1-4* mRNAs have been detected in the CA1 neuropil, suggesting they may be locally translated (109, 110). Dendrite-targeted *KCNIP* mRNAs may also undergo alternative splicing for conditional or spatiotemporal modification of Kv4.2 function. In fact, a KChIP4bL to KChIP4a splicing shift occurs in postmortem brain tissue isolated from Alzheimer’s patients resulting in reduced functional I_A_ (111). It is also possible that KChIPs may confer Ca^2+^ regulation to Kv4.x channel complexes through a mechanism unrelated to direct KChIP-Ca^2+^ binding. KChIP assembly with Kv4.2 is required for downregulation of Kv4.2 surface expression in response to PKA Ser552 phosphorylation (112). Additionally, reduced Kv4.x function by arachidonic acid is also mediated by assembly with KChIPs (113). Thus, there may be ultrastructural roles of KChIPs that augment Kv4.2 in such a way as to affect sensitivity to various forms of regulation.

The majority of CA1 dendritic I_A_ is mediated by Kv4.2 subunits, ruling out a significant contribution of other A-type channels that include Kv1.4, Kv3.4 and Kv4.x subtypes (13). Here, we show that the I_A_ gradient in dendrites is disrupted in the Cav2.3 KO mouse (**Figure 5C**). Others have shown that Ni^2+^-sensitive VGCCs are a source of Ca^2+^ in dendrites and spines (9, 56, 78, 79). One hypothesis, given the results presented here, is that tonic Cav2.3 activity in distal dendrites sustains Kv4.2 functional expression. The increasing distal dendritic gradient of Kv4.2 expression correlates with the ratio of excitatory and inhibitory synapses along the apical dendrite layers of the hippocampus (114, 115). Therefore, ongoing spontaneous excitatory synaptic transmission and Cav2.3-mediated Ca^2+^ entry may maintain Kv4.2 expression as a homeostatic mechanism to regulate local dendritic excitability. Long-term potentiation (LTP) of synaptic inputs may override this by stimulated endocytosis of Kv4.2 in an NMDAR-dependent mechanism as we have previously described (63, 83). This would fit with the reported increase in excitability of dendritic segments containing potentiated synapses following Shaffer Collateral- CA1 LTP (94). Our findings suggest several avenues of research into the function of the Cav2.3-Kv4.2 complex in dendrite function and plasticity; however, a lack of Cav2.3 specific pharmacology without overlapping effects at Kv4.x channels makes this work challenging. Future studies aimed toward identifying Cav2.3-Kv4.2 interaction domains could be leveraged to disrupt the complex to isolate complex-dependent functions.

## STAR METHODS

### Mammalian expression vectors

Cav2.3-GFP was a generous gift from Ehud Isacoff, University of California Berkeley (116). The SGFP2-N1 plasmid was generated by replacement of YFP in YFP-N1 using AgeI/BsrGI sites to excise SGFP2 from pSGFP2-C1 (Dorus Gadella, Addgene 22881 (117)). Kv4.2-CFP, Kv4.2- YFP, and Kv4.2-SGFP2 fusions were created by ligating the mouse Kv4.2 (CCDS29974.1) from Kv4.2-GFP (83) into the CFP-N1 vector using SalI/BglII sites. YFP-Cav2.3 was generated by PCR amplification of the human Cav2.3 (CCDS55664.1) coding sequence from Cav2.3-GFP using BglII/HindIII sites for ligation into the YFP-C1 vector (performed by Bioinnovatise, Inc). Rat pCMV-KChIP2c was generously provided by Henry Jerng and Paul Pfaffinger, Baylor College of Medicine, Houston, TX. Ca^2+^-dead KChIP2c was generated by site directed mutagenesis using D->A mutations at position 1 of each of EF2,3, and 4 (Stratagene, QuikChange Site-Directed Mutagenesis Kit). KChIP2c-CFP was generated by PCR amplification of the rat KChIP2c ORF (NM_001033961.1) from pCMV-KChIP2c and subcloned into CFP-N1 using BglII/SalI sites. AKAP79-YFP and PKARII-CFP were gifts from Mark L. Dell’Acqua, University of Colorado School of Medicine, Aurora, CO. The CFP-18aa-YFP tandem fusion construct used for FRET efficiency calibrations was a gift from Clemens Kaminski (University of Cambridge). GCaMP6f was a gift from Douglas Kim & GENIE project (Addgene 40755 (118)). Human Kv4.2-Myc-DDK (Origene, RC215266), ECFP-N1, EYFP-C1, EYFP-N1, and mCherry-N1 are commercially available (Takara Bio).

### Antibodies

*Guinea pig anti-Cav2.3* was a generous gift from Akos Kulik, University of Freiburg (52), 1:100 for EM; 1:1000 for IHC; 1:5000 for WB. *Mouse anti-Kv4.2* (K57/1): NeuroMab 75-016, 1:25 for EM; 1:300 for IHC; 1:2000 for WB. *Mouse anti-Myc*, Millipore 05-419, 1:500 for ICC. *Alexa Fluor 488 goat anti-guinea pig*: ThermoFisher A11073, 1:800 for IHC. *Alexa Fluor 488 goat anti-rabbit*: ThermoFisher A11008, 1:500 for ICC. *Alexa Fluor 555 goat anti-mouse*: ThermoFisher A21422, 1:800 for IHC. *Alexa Fluor 647 goat anti-mouse*: ThermoFisher A21236, 1:500 for ICC. *Alexa Fluor 680 goat anti-mouse*: ThermoFisher A21057, 1:10,000 for WB. *IRDye 800CW goat anti-rabbit*: Li-Cor Biosciences 926-32211, 1:5,000 for WB. *Rabbit anti-GFP*, ThermoFisher A11122, 1:500 for ICC; 1:3000 for WB

### Cell culture

HEK293FT and COS7 cells were maintained in DMEM supplemented with 10% fetal bovine serum (ThermoFisher, A3160501) and 2% penicillin/streptomycin (ThermoFisher, 15140122) at 37°C and 5.0% CO_2_. Cells were passaged 2x weekly by seeding 0.5-1.0 x 10^6^ cells into 10 cm culture dishes (Corning). Cell lines were kept up to passage 20.

### Humane rodent care and use

All protocols and procedures were approved by the National Institute for Child Health and Human Development Animal Care and Use Committee. All mice were housed and bred in the Porter Neuroscience Research Center animal facility at the National Institutes of Health in Bethesda, MD. Rodents were maintained on a 12 h light/dark cycle with *ad libitum* access to rodent chow and water. Cav2.3 KO mice used for hippocampal cultures and brain slice electrophysiology were generously provided by Dr. Richard Miller, Northwestern University (61). Cav2.3 KOs were maintained on a C57Bl/6J background. Age-matched wild-type C57Bl/6J mice (WT) were used as controls. Rat hippocampal neuronal cultures were prepared with embryos collected from E18-19 timed pregnant Sprague Dawley rats bred at Taconic Biosciences and housed at the NIH for 4-5 days prior to euthanasia.

### Primary culture of rodent hippocampal neurons

Neuronal hippocampal cultures prepared from embryonic day 18-19 (E18) rodent embryos were performed as reported previously (119). Female dams were euthanized using CO_2_ asphyxiation followed by guillotine decapitation. Embryos were rapidly dissected from the uterine horn, decapitated with sharp scissors, and whole heads were placed in ice-cold dissection medium (ThermoFisher, 1X HBSS (14185052), 1 mM sodium pyruvate (11360070), 10 mM HEPES (15630080), and 30 mM Glucose). After peeling away overlying skin and bone with forceps, brains were removed from the skull and placed into fresh dissection medium. Each hemisphere of the cerebral cortex was bisected from the hindbrain and the hippocampus was then gently rolled away and excised from the cerebral cortex and placed into fresh ice-cold dissection medium. Once all tissue was collected, the dissection medium was replaced with 5 ml room temperature (RT) papain solution (5 ml dissection solution w/ 1% DNase (Worthington, LK003170) and 1 vial 0.22 μm filtered Papain (Worthington, LK003176). After a 45 min RT incubation, tissue was washed 4x with prewarmed NB5 medium (5% FBS (Hyclone, SH30071.03), ThermoFisher: 1X Neurobasal A (21103049), 2% Glutamax (35050061), and 2% B27 (17504044). Tissue was dissociated by gentle trituration using a 5 ml plastic serological pipette, cells were filtered through a 70 µm cell strainer (Corning, 352350) and pelleted at 1,000 rpm in a swinging bucket centrifuge (Beckman Coulter Allegra^TM^ 6R) for 5 min at RT. Cells were resuspended in 10 ml NB5, diluted 1:1 in 0.4% Trypan Blue Stain (ThermoFisher, 15250061) and counted with a hemocytometer. Neurons were plated 125,000 (rat) or 175,000 (mouse)/well in a 12–well plate (Corning) containing glass coverslips. 12 mm round Poly-D-lysine/laminin pre-coated glass coverslips (Corning, 354087) were used for electrophysiology. For immunostaining and fluorescence microscopy neurons were plated on in-house Poly-D-lysine/laminin coated 18 mm German glass coverslips (Carolina Biological, 633013). Briefly, UV-sterilized coverslips were incubated overnight in poly-D-lysine (Sigma, P7280-5MG) dissolved in 22 µm filter-sterilized borate buffer (50 mM boric acid, 12.5 mM sodium borate, pH 8.5). The following day, coverslips were washed using sterile water and coated with 0.01 mg/ml mouse Laminin (ThermoFisher, 23017015) in PBS for 3 hrs. 24 hrs after seeding, NB5 was replaced with Neurobasal A (Invitrogen) supplemented as above but without FBS and 1% Glutamax (NB0). Neurons were fed twice per week thereafter by replacing 0.4 ml with 0.5 ml fresh NB0 per well.

### Hippocampus area CA1 double immunogold electron microscopy

Animals used for postembedding, double-immunogold localization were prepared as described previously (120). Briefly, two male, adult Sprague Dawley rats were perfused with phosphate buffer, followed by perfusion with 4% paraformaldehyde + 0.5% glutaraldehyde in phosphate buffer, and then the brains were vibratomed, cryoprotected in glycerol overnight, frozen in a Leica EM CPC (Vienna, Austria), and embedded in Lowicryl HM-20 resin in a Leica AFS freeze-substitution instrument. Thin sections were incubated in 0.1% sodium borohydride + 50 mM glycine in Tris-buffered saline plus 0.1% Triton X-100 (TBST), then in 10% normal goat serum (NGS) in TBST, and then with 2 primary antibodies together in 1% NGS/TBST (overnight); then they were incubated with the 2 immunogold-conjugated secondary antibodies (5+15 nm; Ted Pella, Redding, CA, USA) in 1% NGS in TBST with 0.5% polyethylene glycol (20,000 MW), and stained with uranyl acetate and lead citrate. Controls on sections from the same two rats, labeled with the 2 secondary antibodies but without the primary antibodies, showed only rare gold and no colocalization.

### Native hippocampal co-immunoprecipitation assays

We performed native co-IP experiments to confirm an interaction between endogenous Cav2.3 and Kv4.2 channel with male, 12-week-old wild type C57BL/6 or control Cav2.3-KO mouse hippocampus. Brain hippocampal tissue were lysed in lysis buffer: 150 mM NaCl, 20 mM Tris-HCl, 1% CHAPS and protease inhibitor mixture (Roche, USA) and incubated for 20 min on ice, then sonicated 5 times for 5 s each. The lysate was centrifuged at 15,000 xg for 20 min at 4°C and supernatants were incubated with anti-Cav2.3 (2 µg/500 µg protein) and guinea-pig IgG (ThermoFisher Scientific) as nonspecific control. The mixture was then incubated and rotated at 4°C overnight. The antibody-antigen complex was adsorbed onto 50 μl of immobilized protein A (ThermoFisher Scientific) and incubated and rotated for 2-3 h at 4 °C. The protein-bead mixtures were washed 5x with lysis buffer. The beads were resuspended in SDS sample buffer (ThermoFisher, NP0007) to elute bound proteins. Protein complexes were immunoblotted as described below.

### COS7 cell surface biotinylation

Biotinylation assays were performed as previously described (83). COS7 cells are our preferred cell line for surface biotinylation because of a higher surface to volume ratio and slower dividing time relative to HEK293 cells. COS7 cells were transfected with Kv4.2 and Cav2.3-GFP constructs using X-tremegene 9 transfection reagent (Sigma-Aldrich, 06365779001) for 24-36 h; the cells were rinsed with ice-cold PBS, and surface protein was biotinylated with 1.5 mg/ml sulfo-NHS-SS-biotin reagent (ThermoFisher) in PBS for 30 min on ice. Unbound biotin was quenched with cold 50 mM glycine in PBS. Cells were lysed with ice-cold lysis buffer: 150 mM NaCl, 20 mM Tris-HCl, 1% CHAPS and protease inhibitor mixture (Roche Diagnostics), sonicated and centrifuged at 12,000 g for 10 min. Cell lysates were incubated overnight at 4°C with immobilized-Streptavidin agarose beads (ThermoFisher), and unbound protein was removed from the beads with 5 washes in lysis buffer. The bound proteins were eluted with SDS sample buffer (ThermoFisher, NP0007). Surface expressed and total proteins were immunoblotted as described below.

### Western blots

Sample proteins were separated on 3-8% Tris-acetate gels (ThermoFisher, EA03752) using SDS buffer (ThermoFisher, NP0002) and electrophoresis chambers (ThermoFisher, El0001). Proteins were transferred using electrophoretic chambers (Bio-Rad, 1703930) in a Tris based buffer (ThermoFisher, NP0006) to PVDF membranes (Millipore, IPFL00010). Immunoblots were blocked for 1 h at RT and probed using primary antibodies overnight at 4°C in Odyssey blocking buffer (Li-Cor Biosciences). After washes, immunoblots were probed for 1 h at RT with secondary antibodies conjugated to infrared dyes for detection using the Odyssey infrared imaging system (LI-COR Biosciences, Lincoln, NE). Quantification of results was performed using Odyssey software.

### Immunostaining

*Brain slices:* Deeply anesthetized mice were transcardially perfused with ice-cold paraformaldehyde (PFA) solution (4% PFA (Electron Microscopy Sciences, 15714-S), PBS, pH 7.4). Whole brains were dissected, post-fixed in 4% PFA for 4 h, and cryopreserved with increasing concentrations of sucrose solutions (10, 20 and 30% for 4, 12, and 12-24 h respectively). Cryopreserved brains were sectioned in a cryostat at 7 µm thickness (Histoserv, Inc.) through the dorsal hippocampus beginning at -1.955 mm caudal to bregma and adhered to microscope slides (Superfrost™ Plus, ThermoFisher, 1255015) and stored at -80°C. After thawing at RT for 15 min, a circle was drawn with a pap pen around each section to create a hydrophobic barrier. Sections were rehydrated using PBS for 5 min at RT and blocked for 1 h at RT (0.3% Triton X-100, 1% normal goat serum, PBS, pH 7.3-7.4). Next, fresh blocking solution containing primary antibodies was applied and incubated 24-48 h at 4°C in a closed, humidified box. Slides were washed 4 x 15 min in PBS with agitation. Secondary antibody was applied in 0.2% Triton X-100 PBS and sections were incubated while protected from light for 2 h at RT. Sections were washed 4 x 15 min in PBS with agitation. Coverslips were mounted using a DAPI-containing mounting medium (ProLong^TM^ Gold Antifade, ThermoFisher, P36931).

*Cultured hippocampal neurons:* Primary neurons grown on glass coverslips were washed 1x with PBS and fixed in PFA solution for 10 min at RT. Neurons were washed 3 x 5 min with PBS and permeabilized using 0.2% Triton X-100 for 10 min at RT. After 3 x 1 min PBS washes, neurons were blocked with 3% BSA dissolved in PBS overnight at 4°C and protected from light. Primary antibodies were applied in 3% BSA PBS for 2 h at RT with gentle agitation. Neurons were washed 3 x 1 min in PBS and secondary antibodies dissolved in PBS were incubated for 1 h at RT with gentle agitation and protected from light. After 3 x 1 min washes, coverslips were mounted face-down on glass slides in mounting medium with DAPI (ProLong^TM^ Gold Antifade, ThermoFisher, P36931).

### Confocal fluorescence microscopy and analysis

PFA-fixed neuronal cultures and mouse brain slices were imaged using a Zeiss 710 laser scanning confocal microscope running Zen Black software (Zeiss Microscopy). Fluorescence was acquired using 405/449 nm (DAPI), 488/515 nm (Alexa 488), and 633/670 (Alexa 647) nm laser excitation/emission wavelengths. Hippocampal sections were imaged using a Zeiss 10x Plan-Apochromat 10x/0.45NA air objective capturing 0.83 μm/pixel in *x* and *y* dimensions. Z-stacks were 20.83 μm tall with a 5.21 μm step size. A Zeiss 63x Plan-Apochromat/1.4NA oil objective was used for cultured neurons yielding 0.13 μm/pixel in *x* and *y* dimensions. Z-stacks were 3.36 μm tall using 0.42 μm z-steps, and max intensity projections were used for analysis. Analysis was performed in ImageJ (NIH). Spine and dendrite fluorescence intensities were measured by masking all clearly identifiable mushroom-shaped spines and adjacent dendrite segments from which they projected. Spine/dendrite ratios were calculated from the mean spine and dendrite shaft intensities from each cell.

### Live-cell FRET microscopy in HEK293FT cells

*FRET Acquisition:* Cultured HEK293FT cells were trypsinized ≤ 2 min, counted, and 75,000 cells were seeded onto uncoated 18 mm glass coverslips. After 24 h growth, cells were transfected using OPTI-MEM serum free medium, X-tremeGene 9 (Sigma, 6365779001) and various plasmids. Living cells were imaged in a Tyrode’s salt solution (in mM: 135 NaCl, 5 KCl, 2 CaCl_2_, 1 MgCl_2_, 25 HEPES, 10 glucose, pH 7.4) at RT 24–48 h post-transfection. An Observer.Z1 microscope (Zeiss) with a 63x plan-apochromat, 1.4 NA oil objective (Zeiss), Lambda LS Xenon Arc Lamp Light Source System (Sutter Instruments), AxioCam MRm camera (Zeiss), and Zen Blue software (Zeiss) were used for image acquisition. Three-filter FRET images were captured using appropriate filter cubes (Semrock) housed in the microscope turret. CFP cube: (Ex. 438/24 nm, Em. 483/32 nm, Di. 458 nm), YFP cube: (Ex. 500/24 nm, Em. 542/27 nm, Di. 520 nm), and FRET cube: (Ex. 438/24 nm, Em. 542/27 nm, Di. 458 nm). ImageJ software (NIH) was used for image processing and calculations of sensitized FRET efficiency were adapted from the method of Clemens Kaminski (121) with more details provided below.

*FRET analysis:* CFP, YFP, and CFP-YFP rawFRET fluorescence were captured in single *xy* planes using the following excitation and detection scheme:

CFP image: CFP excitation and CFP emission (CFP fluorescence intensity)

YFP image: YFP excitation and YFP emission (YFP fluorescence intensity)

rawFRET image: CFP excitation and YFP emission (uncorrected FRET fluorescence intensity)

Fluorescence background was estimated by measuring the mean pixel values for several images captured in a cell-free section of the coverslip on the experimental day. After background subtraction, fluorescence intensity in the rawFRET image was corrected for CFP bleed-through and YFP cross-excitation. A significant percentage of the fluorescent signal in the rawFRET image is not due to FRET, but instead results from spectral crosstalk that must be subtracted. A percentage of fluorescence emission from CFP is present in the YFP emission bandpass of the rawFRET image (CFP bleed-through). Conversely, YFP cross-excitation occurs when CFP excitation bandpass in the rawFRET image leads to direct excitation of YFP. For each FRET pair, CFP bleed-through and YFP cross-excitation in the rawFRET image was measured by expressing either the CFP or YFP construct alone and determining the ratio of fluorescence intensity in the rawFRET image divided by the fluorescence intensity in the CFP (donor emission ratio (DER) or YFP (Acceptor emission ratio (AER)) image across many cells. Each FRET pair yields unique spectral cross-talk that when subtracted from the rawFRET image, generates the true signal due to CFP/YFP FRET, known here as corrected FRET (FRETc). The equation

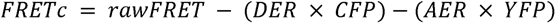

was used to determine the amount of FRET in each cell. Mean CFP, YFP, and raw FRET fluorescence intensities were measured by mask analysis of regions enriched for the construct of interest. For cells expressing 1:1 stoichiometry of CFP and YFP, apparent FRET efficiency values were calculated from mean intensities and normalized to the fraction of acceptor molecules undergoing FRET (*FRET EFF_A_*) using the equation:

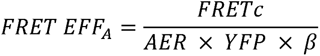

where β is a factor relating spectral and excitation efficiencies of donor and acceptor molecules. For experiments measuring the stoichiometry of CFP/YFP FRET pairs, cells were transfected with various ratios of CFP and YFP cDNAs and imaged as above. In addition to measuring FRET EFF_A_, FRET EFF_D_ was calculated using the equation:

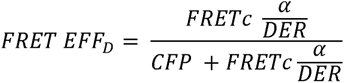

where α relates the quantum yields and signal detection efficiencies between donor and acceptors. The values for α and β were found using the method of Kaminski (121). Briefly, using the wide-field microscopy system described above, transfection of a control CFP-YFP tandem fusion construct consisting of CFP and YFP separated by an 18 amino acid linker (GLRSRAQASNSAVEGSAM) with a predetermined FRET efficiency of 0.38 allows for determination of α and β using the following equations:

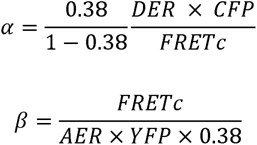

Plotting either FRET EFF_A_ vs. CFP/YFP or FRET EFF_D_ vs. YFP/CFP gives a measure of saturating FRET at either the acceptor or donor. The ratio of the maximum FRET value normalized for acceptor or donor concentration approximates the stoichiometry of the interaction as described by others (68, 122, 123).

### Fluorescence recovery after photobleaching (FRAP) microscopy

*FRAP acquisition:* Fluorescence imaging was performed on a Zeiss 710 laser scanning confocal microscope equipped with a 405-30 nm diode laser, tunable Argon laser, DPSS 561-10 nm laser, and a Zeiss 63x Plan-Apochromat/1.4 NA oil objective. A stage insert incubation system was used to maintain cells at 34°C. Samples were illuminated at low laser power (2.0-3.5%). Image acquisition and ROI bleaching was driven by Zen Black software (Zeiss Microscopy). Images were acquired at 4X zoom yielding 0.033 μm/pixel. Cells were imaged 24-48 hrs after transfection in Tyrode’s salt solution. For HEK293FT cells, 500,000 cells were seeded onto 25 mm coverslips in 6-well cell culture dishes (Corning). After 24 h, cells were transfected with CFP, YFP, Kv4.2-CFP, or YFP-Cav2.3 constructs using X-tremeGene 9 Transfection Reagent (Sigma-Aldrich, 06365779001). Samples were illuminated at low laser power with either 405 nm or 515 nm laser excitation, PMT gain of 700, and at 0.5 Hz. Both CFP and YFP fluorescence were photobleached within a 50 px^2^ ROI using 8 iterations of the 405 nm laser at 100% power. For cultured neurons, DIV12-13 neurons were transfected with Kv4.2-sGFP2 and mCherry using Lipofectamine 2000 Transfection Reagent (ThermoFisher, 52887). Samples were illuminated with 488 nm and 560 nm laser excitation, PMT gain of 700-750, and at 0.2 Hz. Kv4.2-SGFP2 fluorescence was photobleached within a 30 px^2^ ROI for dendrite shafts and a custom ROI matching the shape of each spine. Both dendrite and spine photobleaching required 8 iterations of the 405 nm laser at 100% power.

*FRAP data analysis:* FRAP data was processed and analyzed as previously described (124) using the double normalization method described by Phair and Misteli (125),

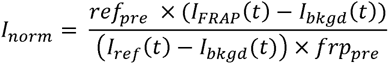

Where *ref_pre_* and *frp_pre_* are the background corrected means of the pre-bleach time points from the reference and FRAP region intensity traces, respectively. *I_FRAP_*(*t*) is the fluorescence intensity within the FRAP ROI, *I_bkgd_*(*t*) is a background intensity from an ROI outside the cell, and *I_ref_*(*t*) is the reference intensity in an unbleached region of the cell to account for photobleaching during acquisition. FRAP curves from individual ROIs were vetted for adequate bleaching (≥ 50% post-bleach intensity compared to pre-bleach intensity) and stability of intensity trace (e.g., traces with distortions due to cellular movement or stage-drift were discarded). After screening, normalized curves were then scaled 0 to 1 and averaged. Standard deviation and standard error of the mean were calculated. Averaged curves were fitted with a single-exponential, *FRAP*(*t*)= A ×(1 – e^-τ×*t*^) where *A* is the mobile fraction using GraphPad Prism software.

### HEK293FT cell whole-cell voltage-clamp recordings

HEK293FT cells were seeded onto 35 mm cell culture dishes at a concentration of 500 x 10^6^ cells per dish. After 16-24 h, cultures were transfected with various plasmids (1-2 μg). To each dish, DNA was first mixed with 300 μl Opti-MEM Reduced Serum Medium (ThermoFisher, 31985070) reduced serum medium. Next, 6 μl X-tremegene 9 DNA Transfection Reagent (Sigma-Aldrich, 06365779001) was added and incubated for 10 min at room temperature before dropwise addition to cultures. On the day of recording (24-48 h after transfection), cultures were trypsinized for ≤ 2 min and seeded at low density onto glass coverslips and allowed to adhere ≥ 1 h. Coverslips were then transferred to a recording chamber and superfused (1-2 ml min^-1^) in 95% O_2_, 5% CO_2_ saturated extracellular solution (in mM: 115 NaCl, 2.5 KCl,1.25 NaH_2_PO_4_, 25 NaHCO_3_, 2 CaCl_2_, 1 MgCl_2_, 25 glucose, pH 7.2-7.3; 300 mOsm/l) at RT. Borosilicate patch electrodes were pulled using a two-stage pipette puller (Narishige PC-10) to a tip resistance of 2.5-4.0 MΩ. Patch electrodes were filled with (in mM): 115 KCl, 10 NaCl, 20 KOH, 10 Hepes, 10 EGTA, and 25 glucose (pH 7.3, 290 mOsm). Maximum voltage-gated K^+^ currents were elicited by voltage steps from a holding potential (−70 mV) to -120mV for 400 ms to relieve Kv4.2 channel inactivation and to +60 mV for 400 ms to maximize channel opening. Inactivation rates were measured by fitting the falling phase of macroscopic currents with a double exponential decay. Voltage dependence of activation was performed using the same holding and hyperpolarizing steps as above but with a range of intermediate activation potentials (-100, -80, -60, -40, -30, -20, -10, 0, +10, +20, +30, +40, and +60 mV). Voltage-dependence of inactivation was determined using 400 ms conditioning steps from holding to -140, -130, -120, -100, -80, -60, -40, -20, -10, and 0 mV immediately before a 400 ms step to +60 mV. Recovery from inactivation was measured using two 400 ms voltage steps to +60mV separated by various intervals (5, 10, 15, 20, 25, 50, 100, 200, and 500 ms).

### Cultured neuron whole-cell voltage-clamp recordings

Primary hippocampal cultured neurons were grown on 12 mm coverslips to DIV6-9 for whole cell recordings or DIV21-27 for miniature excitatory postsynaptic current recordings. At the time of recording, a coverslip was transferred from a 12-well culture plate to the recording chamber and superfused in extracellular solution as described above for HEK293FT cells. Patch electrodes were pulled as described above to a tip resistance of 5-7 MΩ and back-filled with an internal solution containing (in mM): 20 KCl, 125 K-gluconate, 5 EGTA, 4 NaCl, 4 Mg^2+^-ATP, 0.3 Na-GTP, 10 HEPES, and 10 phosphocreatine (pH 7.2, 290 mOsm). Once whole-cell configuration was achieved, neurons were held at -60 mV between voltage protocols. I_A_ was evoked with a 400 ms conditioning step to -80 mV to relieve inactivation before stepping to 0 mV for 400 ms. I_A_ was recorded in the presence of bath applied drugs to block contaminating postsynaptic currents: (in μM) 0.5 Tetrodotoxin (TTX) (Tocris, 1069), 1.0 SR 95531 hydrobromide (Gabazine) (Tocris, 1262), 10.0 CNQX-Na^2^ (Tocris, 1045), and 2.0 MK-801 maleate (Tocris, 0924) or 50 μM D-AP5 (Tocris, 0106). The voltage protocol was repeated every 5 seconds for 6 min. Drugs were applied by rapid perfusion (3 ml/min) 20 sec after recordings began. IC_50_ for Ni^2+^ block of I_A_ was determined using a single exponential fit to the dose-response curve. For measurements of the voltage-dependence of inactivation, 400 ms conditioning steps from holding to -120, -100, -80, -70, -60, -50, -40, -20, -10, and 0 mV immediately before a 400 ms step to 0 mV. Inactivation curves were fit to a Boltzmann function. Miniature excitatory postsynaptic currents (mEPSCs) were recorded in the presence of TTX and Gabazine. For mEPSC recordings in the presence of bath applied Ni^2+^ or AmmTX3, a 10 min wash-in period ensured efficient R-current and I_A_ block, respectively before patching. Immediately after break-in, mEPSCs were recorded in 2 min epochs at a holding potential of -70 mV. Event amplitude was measured by the “Threshold Search” event detection procedure in Clampfit. A maximum of 200 events from each neuron were included for analysis.

### Hippocampal slice preparation and CA1 pyramidal neuron recordings

6-8 week-old male mice were anesthetized with isoflurane and decapitated. For cell-attached dendritic recordings, mice were trans-cardially perfused with ice-cold slicing solution before decapitation. Brains were then transferred into ice-cold slicing solution containing in mM: 2.5 KCl, 28 NaHCO_3_, 1.25 NaH_2_PO_4_, 7 Glucose, 0.5 CaCl_2_, 7 MgCl_2_, 233 Sucrose, and bubbled with 95% O2 / 5% CO2. Transverse slices of the hippocampus (300 µm) were made using a Leica VT1200S vibrating microtome. Slices were transferred to 32°C ACSF containing in mM: 125 NaCl, 2.5 KCl, 25 NaHCO_3_, 1.25 NaH_2_PO_4_, 25 Glucose, 2CaCl_2_, 1 MgCl_2_, 1 ascorbic acid, 3 Na-Pyruvate, and bubbled with 95% O2 / 5% CO2. After 25 minutes the slice chamber was transferred to room temperature for the rest of the recording day. For recording, slices were transferred to a recording-chamber with continuous flow of ACSF (2-3ml/min). Recording pipettes for somatic whole cell and outside-out patch recordings had a tip-resistance of 3-5 MΩ and were filled with an internal solution containing in mM: 20 KCl, 125 K-Gluconate, 1 EGTA, NaCl, 4 NaCl, Mg-ATP, 0.3 NaGTP, 10 HEPES, 10 Phosphocreatine, pH 7.26, 296 mOsm. Recording pipettes for dendritic cell-attached voltage-clamp were filled with an internal solution mimicking ACSF, containing in mM: 3 KCl, 125 NaCl, 10 HEPES, 25 Glucose, 2CaCl_2_, 1 MgCl_2_, pH 7.24, 290 mOsm. Dendritic cell-attached recordings were performed in voltage-clamp mode and voltage step protocols have been inverted for readability in the results section, to account for the reversed polarity of the cell-attached recordings. Protocols were designed for a theoretical resting membrane potential of -60mV. I_A_ was calculated by subtracting I_sus_ from I_tot_ as previously described (59). I_tot_ was elicited using a voltage step to +40 from a -120 mV pre-pulse and a subsequent step to +40 from -30 mV was used to isolate I_sus_. Bath ACSF was supplemented with 1µM TTX to block voltage-gated sodium channels, 1µM Gabazine to selectively block GABAA receptors, and 10µM CNQX to block AMPA-type and Kainate-type glutamate receptors. Recording electrodes were pulled to 10-12 MΩ using a Narishige PC-10 pipette puller and polished using a Narishige microforge.

### Electrophysiology equipment, data processing, and analysis

Manual patch clamp experiments were performed on an Axioskop 2 FS Plus microscope (Zeiss) with CP-Achromat 10× 0.25 air (Zeiss) and LumplanFL 60× 1.0 NA water immersion (Olympus) objectives, and Sutter MPC-200 multi manipulator system with ROE-200 controller. The rig was equipped with a Sutter LB-LS/17 lamp for fluorescence and DIC optics. An Axon Multiclamp 700B amplifier, Axon Digidata 1440A A/D converter, and PClamp software (Molecular, Devices, Sunnyvale, CA) were used to acquire electrophysiological signals. Currents were normalized to cell size using whole-cell capacitance upon cell break-in, and leak currents were subtracted using a P/4 protocol. Data were analyzed using Microsoft Excel, MATLAB, IGOR Pro (WaveMetrics, Lake Oswego, OR), and GraphPad Prism. Pooled data are presented as either bar graphs ± SEM overlaid with individual data points or in tabular format ± SEM.

### Cultured hippocampal neuron Ca^2+^ imaging and analysis

Fast confocal imaging was performed using a 25× 1.1 NA water immersion objective on an A1R MP HD system (Nikon) coupled to a Retiga ELECTRO CCD camera (QImaging) used for sample scanning. Time-series were captured at 15.3 Hz using a 6x zoom on a resonant scanner yielding 0.08 μm/pixel. Image acquisition was controlled by Elements software (Nikon) and analysis was performed in ImageJ (NIH). DIV13-15 primary hippocampal neurons grown on 18 mm glass coverslips were transfected 24-48 h before imaging using Lipofectamine 2000. Neurons were transfected with the genetically encoded Ca^2+^ indicator GCaMP6f and mCherry as a marker of neuronal morphology. Coverslips were transferred to a quick release magnetic imaging chamber (Warner, QR-41LP) with constant perfusion (1-2 ml/min) of modified Tyrode’s salt solution (3 mM CaCl_2_, 0 mM MgCl_2_, 10 μM glycine, 0.5 μM TTX) at RT. Variability and low event frequency required identification of spines that were active both before and after pharmacological treatments. Spine Ca^2+^ signals were analyzed by drawing a mask over the spine of interest. Mean GCaMP6f fluorescence intensity was measured using the “Plot Z-axis Profile” function in imageJ. Extracted intensity values were background-subtracted and normalized to baseline fluorescence intensity (ΔF/F_0_). Only a transient > 2.5 fold above baseline was used for analysis. Many of the analyzed Ca^2+^ transients were clearly individual events; however, if two or more transients overlapped, we developed rules to discern whether they should be considered multiple events. For a trailing event to be considered a unique event, the first transient must have dropped >50% of maximum. Additionally, the trailing transient must 50% above the lowest Δ F/F_0_ of the previous spike to be considered a separate event. If a trailing transient did not meet both criteria it was not considered unique from the initiating transient.

## ACKNOWLEDGEMENTS

This work was supported by the Intramural Research Program of the Eunice Kennedy Shriver National Institute of Child Health and Human Development (J.G.M., J.J.G., L.L., J.H., and D.A.H.) and the National Institute of General Medical Sciences Postdoctoral Research Associate Training Grant (FI2 GM12004) (J.G.M.). Support was also provided by the Intramural Research Program of NIH/National Institute on Deafness and Other Communication Disorders (NIDCD) (Advanced Imaging Core-ZIC DC000081) (R.S.P. and Y-X.W.). We thank members of the Hoffman Lab for valuable discussions, protocols, and experimental assistance. We would like to thank Mark Dell’Acqua and for comments on this manuscript. We are grateful to Adriano Bellotti for contributing cultured rat neurons used in some experiments. The Ca^2+^- dead KChIP2c construct was generated by site directed mutagenesis by Jung Park. Ying Liu bred, genotyped, and cared for the WT and Cav2.3 KO mice used in this work. We thank Vincent Schram (NICHD Microscopy & Imaging Core) for technical assistance.

## AUTHOR CONTRIBUTIONS

Conceptualization, J.G.M, J.J.G., J.H. and D.A.H.; Methodology, J.G.M, J.J.G., J.H., and R.S.P.; investigation, J.G.M, J.J.G., L.L., J.H., R.S.P., and Y-X.W.; Writing – Original Draft, J.G.M.; Writing – Review & Editing, J.G.M., J.J.G., L.L., R.S.P., J.H., and D.A.H.; Funding Acquisition, J.G.M. and D.A.H. Supervision, D.A.H.

---

**Supplemental Figure 1 – Related to Figure 1.**
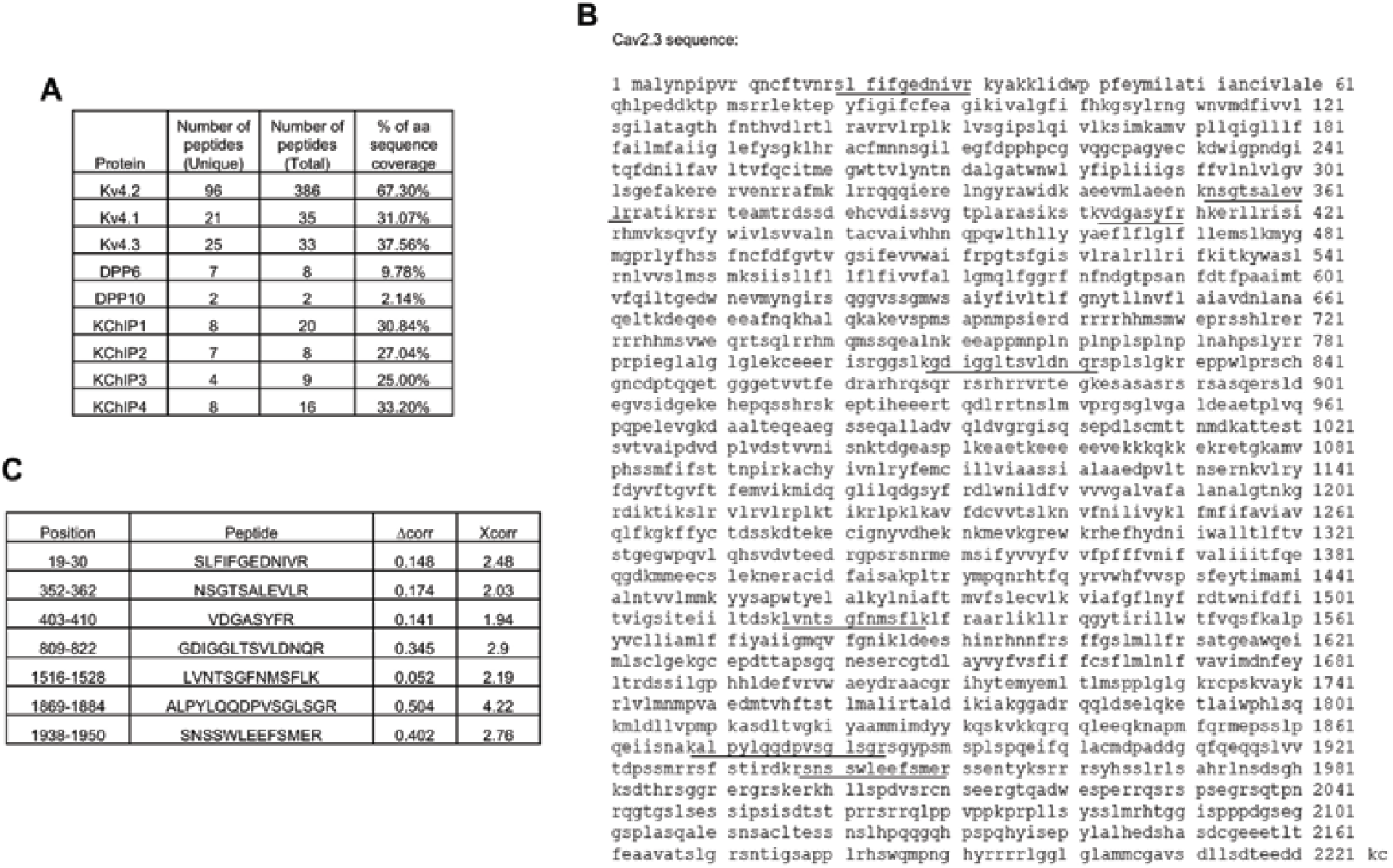
A tandem affinity purification-mass spectrometry screen identified Cav2.3 as a Kv4.2 binding partner. **A.** The numbers of unique and total peptides in Kv4.2 TAP-MS experiments and the percent (%) amino acid sequence coverage for each protein. **B.** Amino acid sequence coverage obtained for Cav2.3 protein from Kv4.2 TAP-MS in rat hippocampal neurons. Peptides detected by MS are underlined. C. Cav2.3 protein sequence report. Xcorr: Sequest cross-correlation score; Δcorr: Xcorr difference between the top ranked and next best sequence.

**Supplemental Figure 2 – related to Figure 1.**
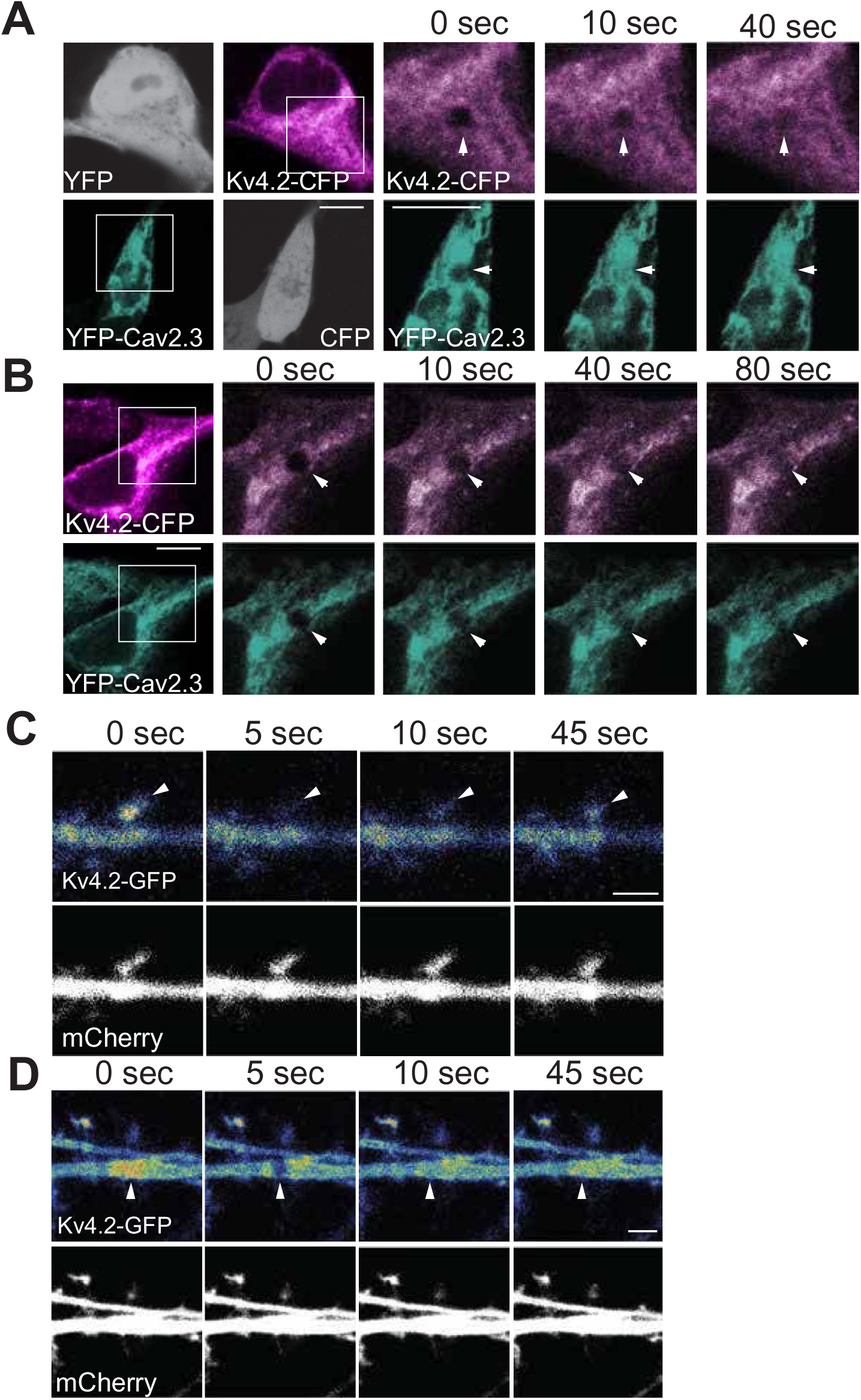
Cav2.3 regulates Kv4.2 mobility in HEK293 cells and neuronal dendrites. **A.** HEK293 cells were transfected with either YFP and Kv4.2-CFP (above) or YFP-Cav2.3 and CFP (below). FRAP time series for Kv4.2-CFP (above) or YFP-Cav2.3 (below) shown in pseudocolor that corresponds with the color coding in Figure 1G. 5 μm scale bars. **B.** Representative images of coexpressed Kv4.2-CFP (above) and YFP-Cav2.3 (below). The simultaneous FRAP timelapses are shown using color coding corresponding to Figure 1H. 5 μm scale bars. **C.** Spine FRAP in a WT mouse hippocampal neuron expressing Kv4.2-SGFP 2 (temperature spectrum pseudocolor) and mCherry (white). 2 μm scale bar. **D.** Dendrite Kv4.2- SGFP2 FRAP is shown as in F. 2 μm scale bar. Arrowheads indicate laser photobleach position.

**Supplemental Figure 3 – Related to Figure 5.**
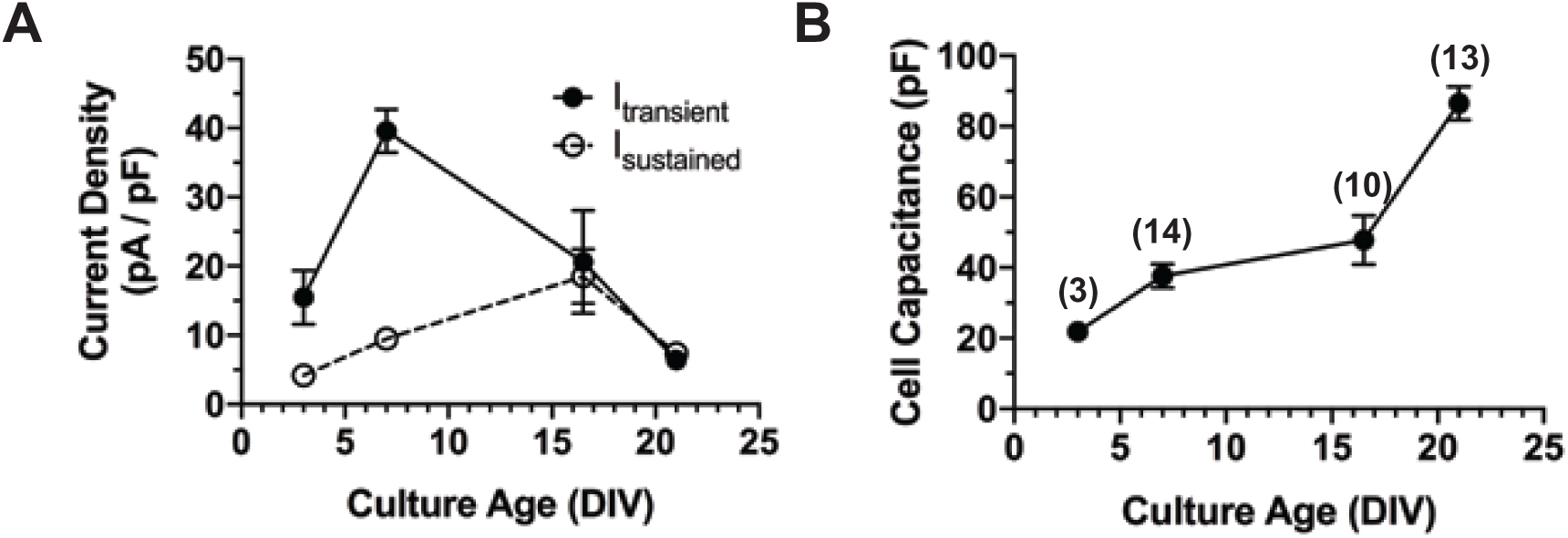
Whole-cell I_A_ predominates in early hippocampal neuron development. **A.** Mean isolated transient (I_A_) and sustained current density plotted by neuronal age in culture. **B.** Cell size measured by whole-cell capacitance on break-in is plotted by neuronal age in culture. Number of cells are indicated in parentheses in **B** but also apply for data points in **A**. Error bars represent +/- SEM from 1-2 cultures.

**Supplemental Figure 4 – Related to Figure 5.**
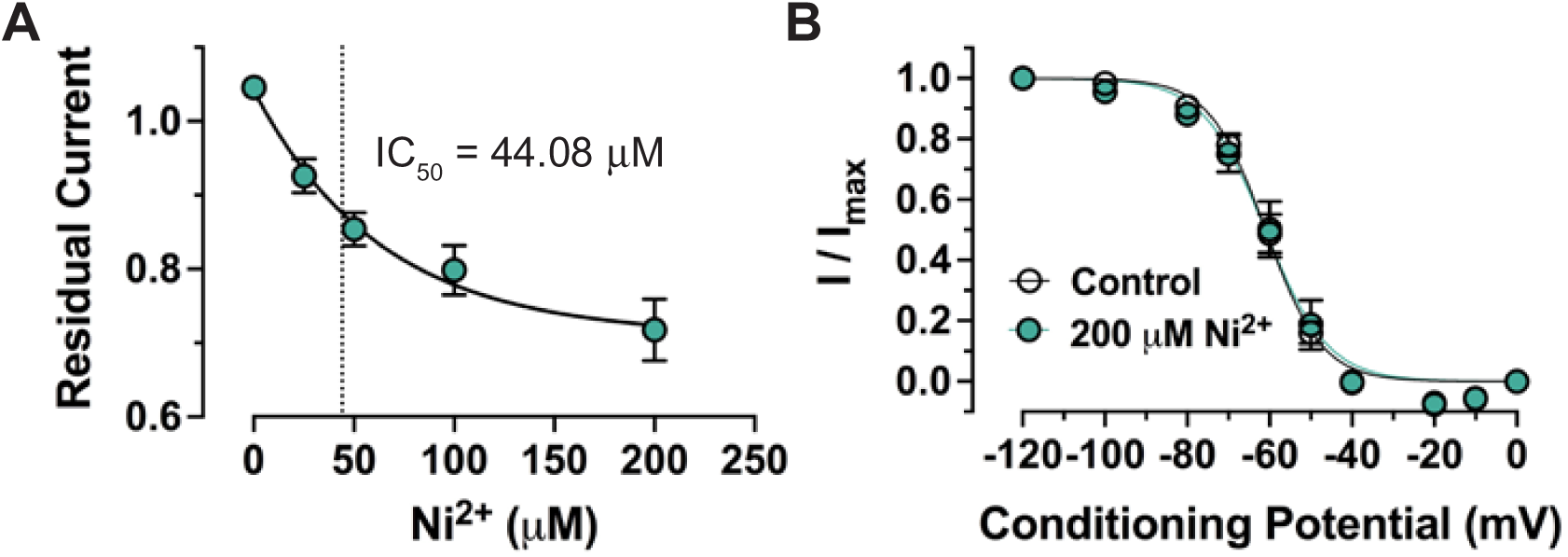
Dose dependent Ni^2+^ inhibition of I_A_ without a shift in voltage-dependence of inactivation. **A.** Residual current was plotted as a function of Ni^2+^ concentration and fit to a single exponential function. The dotted line indicates half-maximal effect of Ni^2+^**. B.** Voltage-dependence of inactivation was determined by holding neurons at a range of conditioning potentials (x-axis) prior to a test potential (0 mV) to determine the effect of Ni^2+^ on the degree of channel inactivation. Data were fit to a Boltzman function. Error bars represent +/- SEM.

**Supplemental Figure 5 – Related to figure 6.**
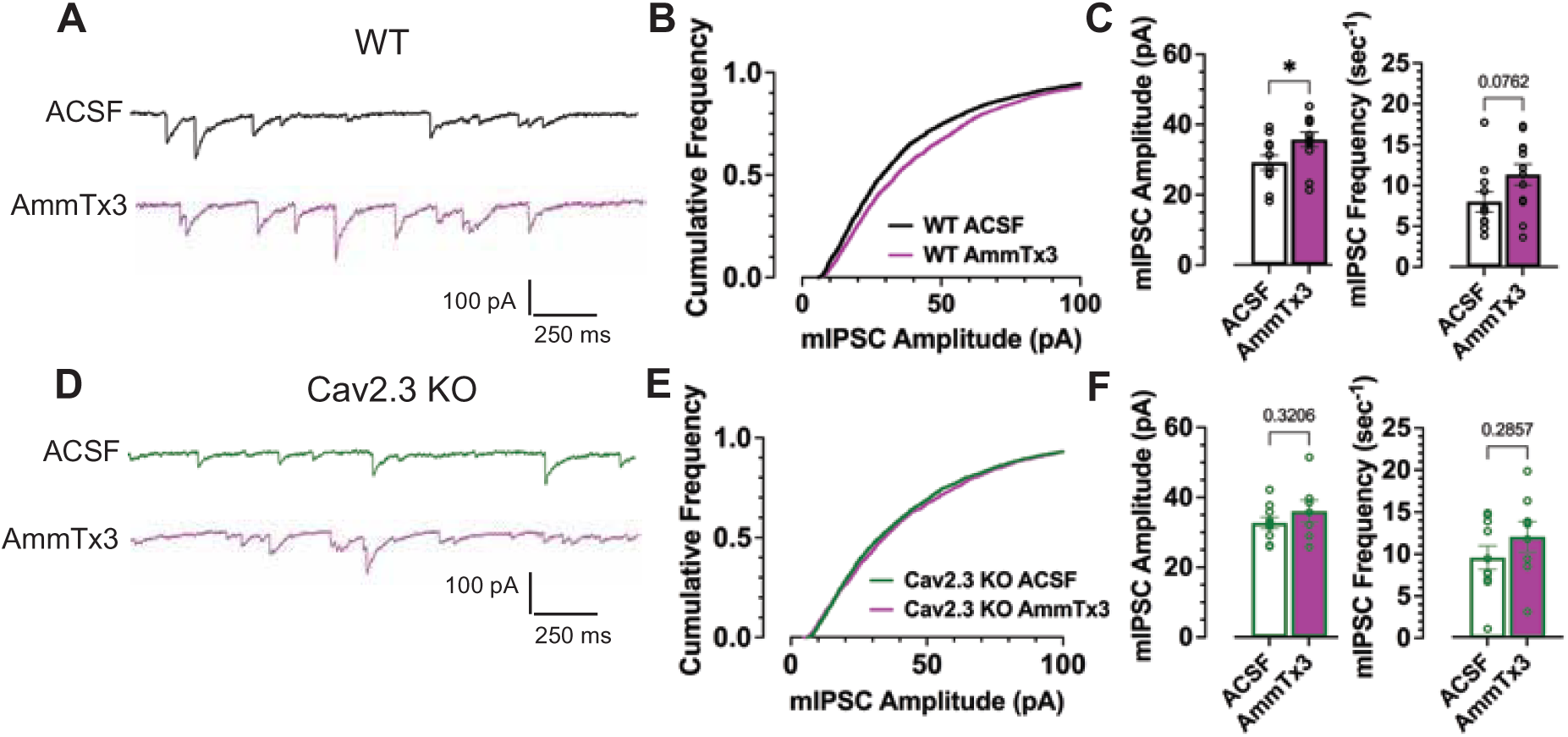
Cav2.3 promotes I_A_-mediated attenuation of spontaneous miniature inhibitory postsynaptic currents. A. Representative mIPSCs recorded from cultured WT mouse hippocampal neurons at a holding potential of -70 mV in the absence (black trace) or presence of AmmTx3 (purple). B. The cumulative distribution of mIPSCs amplitudes demonstrated an AmmTx3-mediated rightward shift in mIPSC amplitudes. C. Median event amplitude (left) and mean frequency (right) from each neuron are plotted. AmmTx3 increases median amplitude. D-F. Cav2.3 KO mIPSCs recorded and summarized as in A-C. AmmTx3 had no effect on mIPSC amplitude in Cav2.3 KO neurons. Error bars represent mean +/- SEM. * *p* < 0.05. Statistical significance was evaluated by unpaired t-test.

